# Multivalent lamin binding controls the meshwork structure in a self-assembled model of the nuclear lamina

**DOI:** 10.64898/2026.03.14.711786

**Authors:** Hadiya Abdul Hameed, Ata Utku Ozkan, Aykut Erbas

## Abstract

The nuclear lamina is a specialized two-dimensional filamentous polymer meshwork that provides structural integrity and elasticity to the nucleus while orchestrating diverse cellular processes. Composed of interacting A- and B-type lamin networks, this structure undergoes tightly regulated self-assembly that is frequently perturbed by disease-causing mutations such as those observed in laminopathies or cardiomyopathies. However, because filament assembly, peripheral adsorption of lamins, and network branching occur concurrently *in vivo*, isolating the specific biophysical parameters that dictate emergent lamina topology has remained a major challenge. Here, we present a polymer-physics approach that explicitly resolves the spontaneous self-assembly of lamin networks under nuclear confinement. By modeling lamin dimers as semiflexible filaments with distinct interactive domains, we demonstrate that the formation of continuous, high-aspect-ratio fibers strictly requires a coordination cascade of parallel lateral alignment sites and longitudinal head-to-tail interactions between lamins. We show that the thermodynamic affinity between lamin-A and the peripheral boundary (i.e., the inner nuclear membrane and lamin B network) acts as a kinetic switch: weak surface adsorption drives network phase separation and large lamin-free gaps, whereas robust substrate binding stabilizes highly branched networks with uniform lamin distribution. Finally, uniaxial compression simulations reveal that mutations altering these molecular binding interfaces severely compromise macroscale nuclear load-bearing capacity and induce structural vulnerabilities. Collectively, our model establishes a predictive, multi-scale view that directly bridges nanoscale lamin interactions with mesoscale topological remodeling of lamina and macroscale nuclear mechanopathology.

## I. INTRODUCTION

The eukaryotic nuclear lamina is a protein meshwork located at the nuclear periphery, positioned between the inner nuclear membrane (INM) and the underlying chromatin. Mutations in proteins that constitute the lamina give rise to a diverse group of diseases collectively known as laminopathies, which include muscular dystrophies, cardiomyopathies, lipodystrophies, and premature aging syndromes [1–7]. The lamina in these disease cells exhibits pronounced morphological abnormalities, including surface holes, wrinkling, and lamina thickening [8– 13]. These structural perturbations compromise lamina integrity and promote nuclear envelope rupture and blebbing, often causing DNA breaks and cell death under cytoskeletal stress [14–17]. However, the precise biophysical mechanisms by which mutations remodel the lamina and affect nuclear mechanics remain elusive.

The nuclear lamina is primarily composed of lamins, a family of type V intermediate filament proteins that assemble into filamentous networks underlying the nuclear envelope [18–20]. Each lamin monomer comprises an intrinsically disordered N-terminal head, a central *α*-helical rod, and a globular C-terminal tail domain [21– 23]. Lamin monomers associate via their *α*-helical rod domains to form parallel coiled-coil dimers *in vitro* [18] (Fig. 1a). These dimers exhibit a conformational behavior reminiscent of semiflexible polymers on surfaces, with a persistence length of *∼* 10 nm [18, 23]. Dimers further assemble into higher-order structures, including protofilaments and thicker fibers with diameters ranging from nm or thicker [19, 22–25]. Cryo-electron tomography studies have revealed network junctions in which four to five lamin filaments converge, forming nodes with a characteristic inter-node spacing around 100 nm [26, 27], albeit with various topologies [12]. Collectively, these observations support a physical description of the nuclear lamina as a network of semiflexible polymers, whose topology and connectivity are likely to be altered under pathological conditions, similar to a polymer gel.

**FIG. 1.**
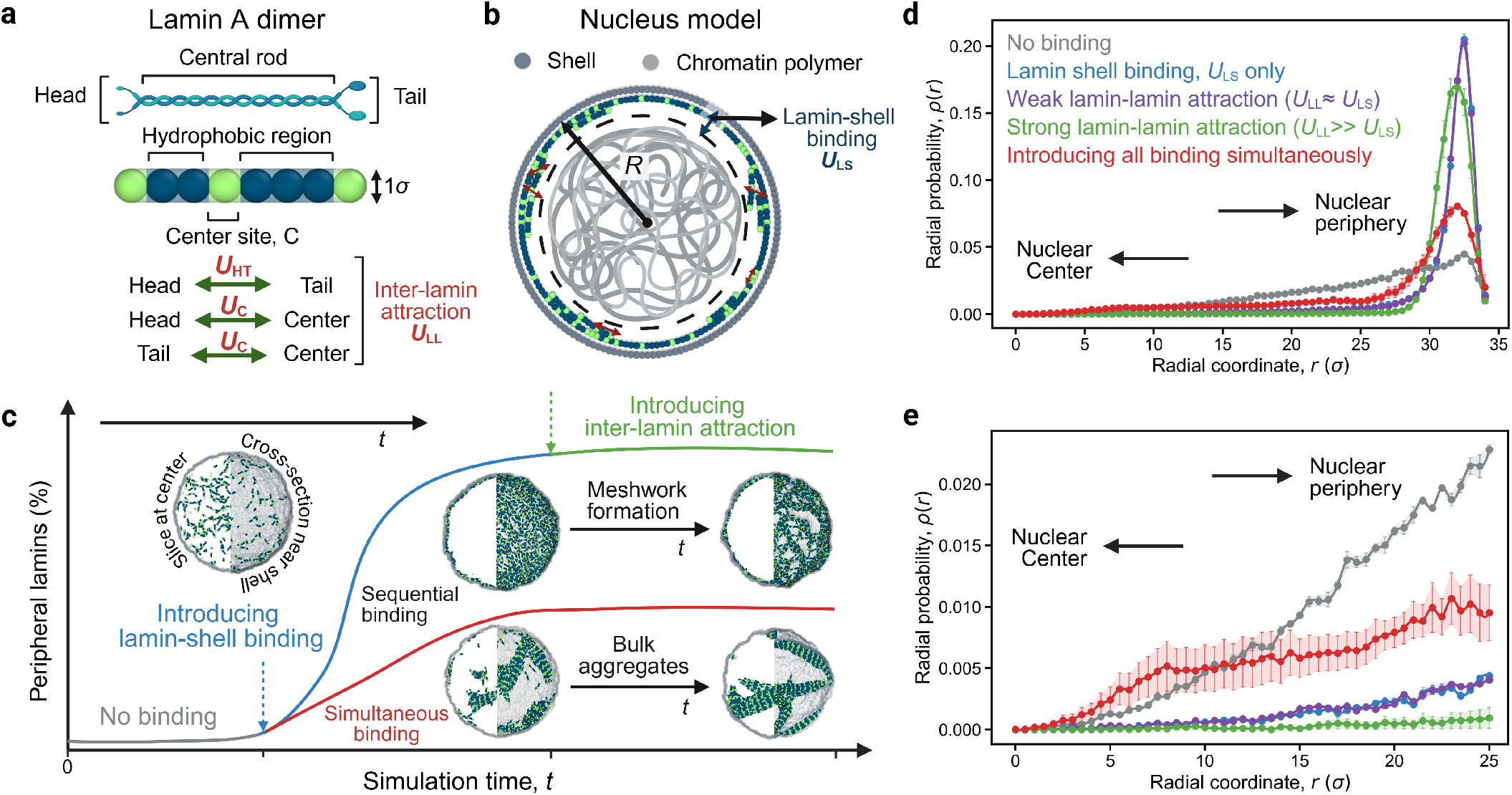
Schematics of the simulation model of lamina self-assembly within the elastic nucleus. (a) Coarse-grained LA dimer. The head, terminal, and additional bead in the central domain are allowed to associate with one another asymmetrically, representing inter-lamin attraction, *U*_LL_. (b) Schematic of the nucleus model composed of an elastic nuclear shell, representing LB and the nuclear boundary, together with an enclosed chromatin polymer (gray chain). Lamin dimers bind to the elastic shell via an affinity, *U*_LS_. (c) Schematic representation of the degree of peripheral lamins with simulation time. Lamin dimers are initially placed randomly within the elastic nucleus (gray), followed by localization where *U*_LS_ is introduced (light blue), and lamina assembly via *U*_LL_ (green). Peripheral lamins decline when both these interactions are introduced simultaneously (red). (d) Radial density, *ρ*(*r*), profiles for various attraction cascades in (c). (e) The same density profile focuses on the interior of the nucleus, highlighting the formation of bulk aggregates when lamin dimers attract before localization. The inset shows the formation of bulk aggregates. For each simulation snapshot, the left half shows a 5*σ*-wide slice taken through the center, while the right half shows the cross-section along the *z*-axis, within 2.5*σ* of the shell surface.

The lamina meshwork is composed of two major classes of lamin proteins, A/C and B-type lamins, each of which can form structurally distinct yet interconnected networks within the lamina [12, 27, 28]. The B-meshwork predominantly occupies the outer nuclear periphery, closely associated with the INM, whereas the lamin-A network is positioned more internally, interfacing and interacting with underlying chromatin [27, 29, 30]. This spatial segregation is consistent with their biochemical differences in a healthy cell: B-type lamins remain permanently farnesylated, promoting their stable anchorage to the INM, while a fraction of A-type lamins resides in the nucleoplasm and undergoes kinetic exchange with the peripheral lamins [8, 31–33]. Although the precise mechanisms governing interactions between these two networks remain incompletely understood [29, 34], emerging evidence from live-cell imaging and computational analyses indicates that B-type lamins play a critical role in stabilizing the lamin-A meshwork. Consistently, depletion of lamin B or silencing of related genes leads to pronounced distortions in the A meshwork, including the formation of lamin-A-depleted domains at the nuclear periphery across various cell types [1, 10, 12, 22, 28, 35– 37]. These observations suggest that membrane-anchored lamin B regulates lamin-A self-assembly at the nuclear surface, promoting a mechanically robust and spatially homogeneous lamina under physiological conditions [27, 34].

The topology and mesoscale organization of the lamina meshwork at the peripheral surface is also affected by various disease-associated lamin mutations that perturb key interaction domains governing lamin dimerization, higher-order filament assembly, and membrane-tethering stability [2, 8, 9, 28, 38–41]. These mutations are thought to interfere with lamina self-assembly and reduce nuclear mechanical stability [10, 11, 15, 16, 36, 42, 43]. For example, mutations that can distort the head-head or rod-rod interactions, such as *Lmna* E161K, K97E, or N195K, directly affect nuclear mechanics, leading to more frequent ruptures [15]. Studies on the crystal structure of progeria mutant S143F protein suggested the formation of X-shaped disulfide bonds in the central rod domains of progerin [44]. This could be a by-product of an increase in the affinity of an additional attractive site within lamin dimers (Fig. 1) [10]. Collectively, available evidence suggests that laminopathic phenotypes arise from perturbed inter-lamin interactions, leading to aberrant assembly of A- and B-type lamins at the nuclear periphery. These molecular-scale disruptions propagate to mesoscale defects in lamina architecture and compromised nuclear mechanics. However, the underlying biophysical interactions responsible for these assembly defects remain poorly understood.

To elucidate the molecular-scale organization of A-type lamins during lamina self-assembly, we develop a *state-of-the-art* polymer physics model of the nuclear lamina. In our model, lamin A dimers are represented as *crosslinkable* semiflexible polymers that self-assemble within an elastic nuclear shell representing the B-meshwork and membrane (Fig. 1). Our results show that the interplay between lamin interactions gives rise to a spectrum of lamin A meshwork states (Fig. 2), including enlarged mesh faces reminiscent of those observed in lamin B-deficient and disease-associated nuclei (Fig. 3) [11, 12, 28]. Further, binding interactions along the rod domains of lamin dimers are essential for the formation of paracrystalline fibers (Fig. 2 and 3). Peripheral lamin organization also affects the mechanical response of the model nucleus under mechanical compression, supporting the view that lamin mutations disrupting the meshwork can weaken nuclear integrity (Fig. 4) [10, 15, 35, 40, 45, 46]. Together, our results demonstrate that a quantitative biophysical framework can link molecular-scale lamin interactions to emergent lamina topology and nuclear rigidity in healthy and diseased nuclei.

**FIG. 2.**
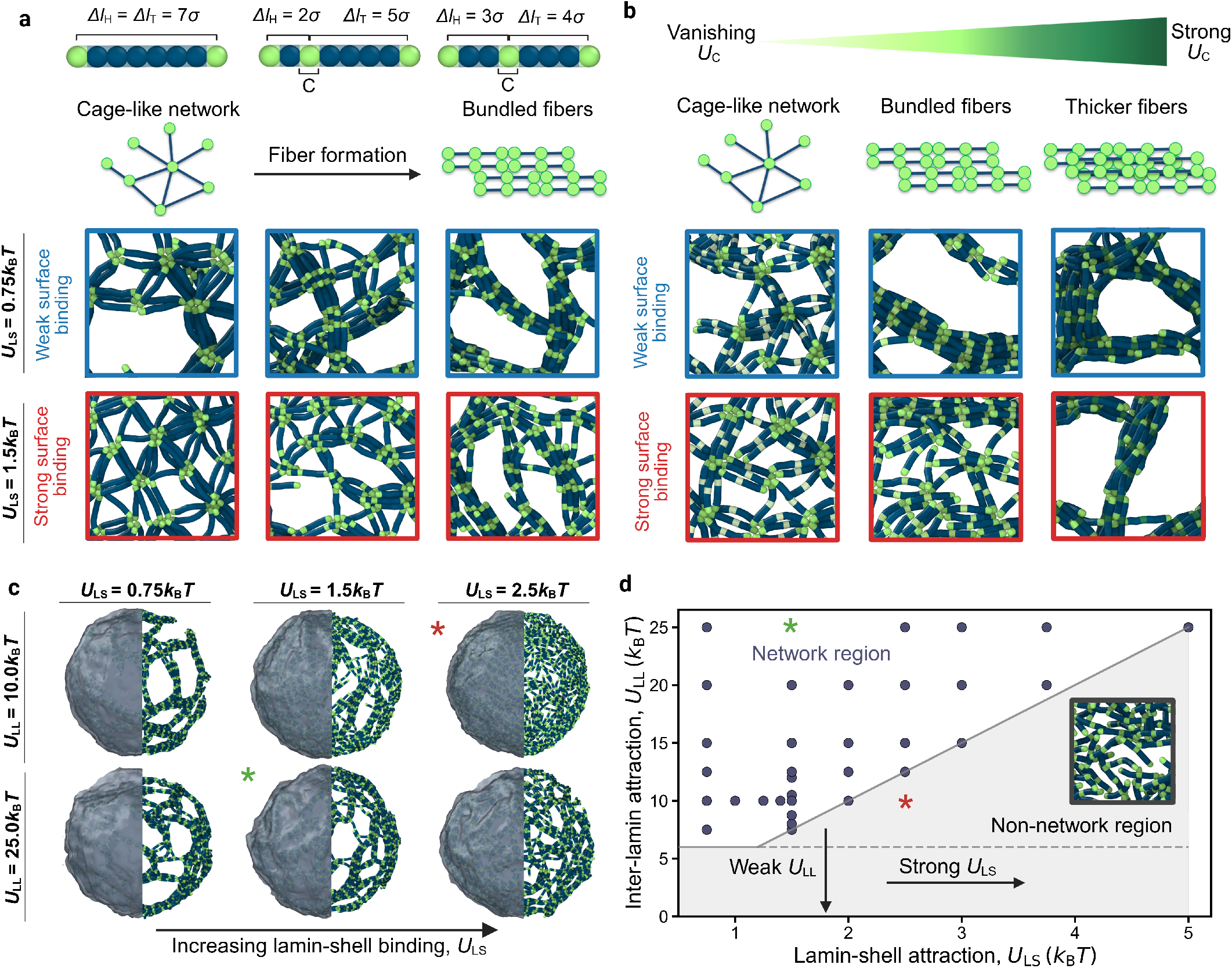
Effect of inter-lamin interaction, *U*_LL_, and lamin-shell binding, *U*_LS_, on lamina meshwork connectivity. (a) Representative lamin network morphologies for different placements of additional attraction site, *C*. Top illustrations indicate the relative positions of the interaction sites along the lamin filament. Introducing an additional attraction leads to a transition from cage-like networks to paracrystalline arrays, under both weak (blue panels) and strong (red panels) lamin-shell binding affinities, *U*_LS_. (b) Representative simulation snapshots illustrating lamina morphologies at increasing central site attraction, *U*_C_. Increasing *U*_C_ promotes thicker paracrystalline fibers, with stronger effects irrespective of lamin-surface binding. (c) Simulation snapshots illustrating the combined effects of lamin–shell binding (*U*_LS_), left to right, and inter-lamin attraction (*U*_LL_), top to bottom, on lamina meshwork connectivity. The left half represents the shell surface, while the right half shows network connectivity. (d) Phase diagram of average node count representing the transition between non-network and network-forming regimes as a function of lamin interactions, *U*_LL_ and *U*_LS_. Asterisks on (c) and (d) represent a network (green) and a non-network (red) configuration.

**FIG. 3.**
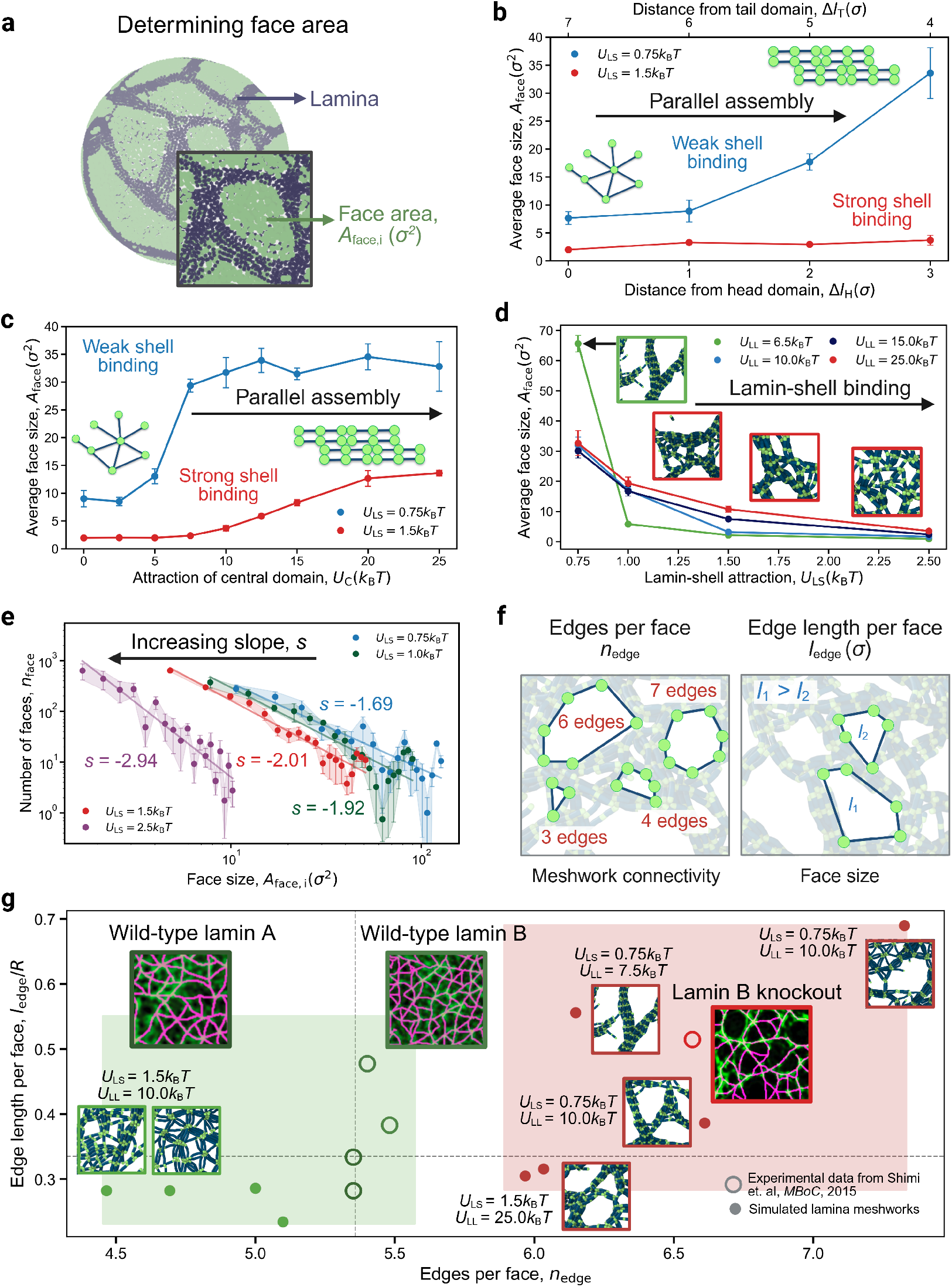
Lamin-free gaps in the lamina meshwork depend on lamin-shell binding affinity, *U*_LS_. (a) Schematic illustrating how lamin-free face area is determined via a geometry-filling procedure. Face area distributions, *A*_face,i_, are extracted to quantify effective mesh pore size [28]. (b) Average face size, *A*_face_, as a function of the separation distance between lamin interaction domains. The bottom axis indicates the distance of the central interaction domain from the head domain, Δ*l*_H_, while the top axis indicates the corresponding distance from the tail domain, Δ*l*_T_. (c) *A*_face_ as a function of central-domain attraction strength, *U*_C_. (d) Average mesh face size, *A*_face_, as a function of lamin-shell binding affinity, *U*_LS_, for increasing inter-lamin attraction strengths, *U*_LL_. Stronger lamin-shell binding produces smaller mesh faces. (e) Distributions of mesh face areas corresponding to the conditions shown in (d), with *U*_LL_ = 25.0*k*_B_*T* . Increasing *U*_LS_ shifts the distributions toward smaller face areas and steeper scaling behavior, consistent with reduced fiber formation. (f) Schematic illustration of the connectivity metrics used to characterize lamina meshworks: the number of edges per face, *n*_edge_, and the average edge length per face, *l*_edge_. (g) Summary of meshwork configurations obtained in (d), shown as *l*_edge_*/R* versus *n*_edge_. Gray lines denote the median values across all simulated conditions. Green squares indicate putative wild-type lamin A cases, whereas red squares indicate *lmnb*^*−/−*^ (lamin-B knockout, lamin-A only) cases. Analogous experimental images and measurements are reported in [12]. In experimental images, green represents m-Emerald-tagged lamin isoforms, while magenta represents the computational reconstruction of the meshwork [12]. Scale bar for experimental images: 1*µ*m.

**FIG. 4.**
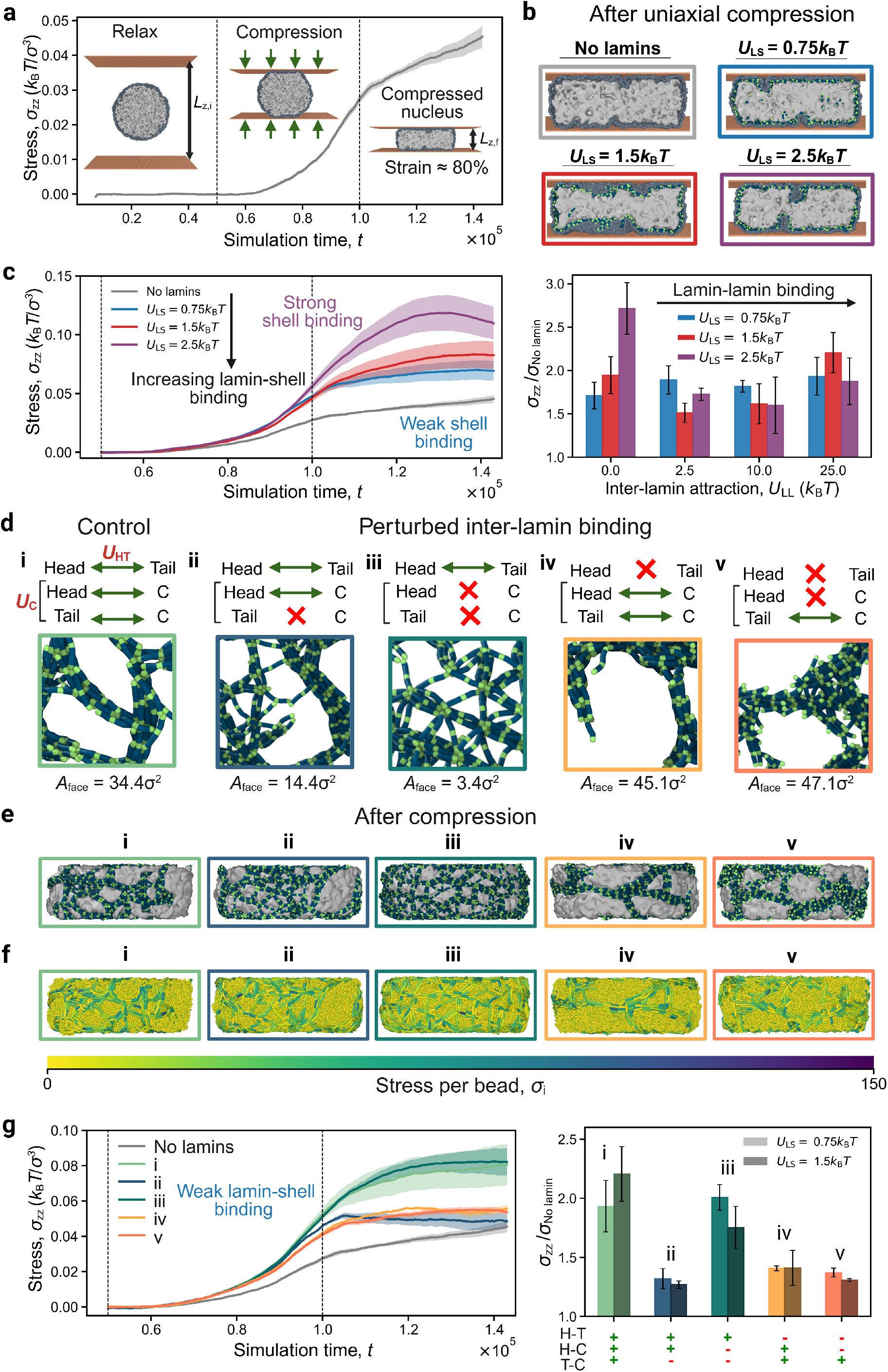
Mechanical response of lamina meshworks under uniaxial nuclear compression depends on lamin-surface binding and perturbations to inter-lamin attraction. (a) Representative stress-time curve in the deformation direction during uniaxial compression on the nucleus. Insets illustrate the compression geometry and progressive nuclear deformation from the initial spherical configuration to the compressed state along the *z*-axis. (b) Representative nuclear cross-sections after uniaxial compression for different lamin-shell binding affinities, *U*_LS_. Colored borders correspond to the simulation conditions shown in panel (c). (c) Left: Stress-time curves for elastic nucleus lacking lamins (gray) and lamina meshworks assembled under increasing lamin-shell attraction only, *U*_LS_. Stronger lamin-shell binding enhances nuclear stiffness and resistance to deformation. Right: Relative maximum stress for stress-time curves, *σ*_zz_*/σ*_No lamin_, as a function of inter-lamin attraction strength, *U*_LL_, for *U*_LS_ in the left panel. (d) Schematic of perturbations to inter-lamin interactions. The control configuration includes attractive *U*_LL_ interactions shown in Fig. 1a. Representative meshwork morphologies and respective gap sizes, *A*_face_, for each perturbation are shown below the schematics. (e) Simulation snapshots of lamina meshworks perturbations, shown in (d), after uniaxial compression. (f) Simulation snapshots in (e) color-coded with stress per bead, *σ*_i_. (g) Left: Stress-time curves for different perturbations of inter-lamin attraction under weak lamin-shell binding, *U*_LS_ = 0.75*k*_B_*T*, shown visually in (d-f). Right: *σ*_zz_*/σ*_No lamin_ for the corresponding perturbation conditions at weak, *U*_LS_ = 0.75*k*_B_*T*, and strong lamin-shell binding, *U*_LS_ = 1.5*k*_B_*T* . Networks with perturbations to the central-domain interaction exhibit softer nuclei. All stress-time curves are smoothed with a moving average.

## II. RESULTS

### A. Peripheral meshwork assembly requires sequential lamin interactions in simulations

Experimental studies have shown that A-type lamins exist as two sub-populations in a healthy cell nucleus. These nucleoplasmic and peripheral populations appear to be rapidly exchanging with one another, indicating that lamin A can interact transiently with itself and its higher-order structures [8, 9, 47, 48]. In fact, nucleoplasmic lamin A/C is enriched in Ser22-phosphorylated forms, keeping it more soluble and mobile, whereas peripheral ones are largely dephosphorylated [47]. In parallel, the lamin-A meshwork protects the nucleus from mechanical damage [15, 45, 49, 50], and thus should be stable enough to withstand mechanical stress. These contradictory requirements point to a self-assembly mechanism in which the formation of the lamin-A meshwork may depend not only on the strength of binding affinities among various lamins, but also on the sequence in which these interactions emerge in space and time.

Hence, we first explicitly test how sequential versus simultaneous activation of lamin-A interactions (with themselves and with the inner surface of the nuclear shell) can govern peripheral lamin localization, aggregation, and ultimately, meshwork formation (Fig. 1c-e). In our mesoscopic nucleus model, a bead-spring elastic shell emulates the B meshwork and INM. This shell provides a steric nuclear confinement and adsorption surface for the peripheral localization of A-type lamins, where each lamin-A dimer is modeled as a short, semiflexible polymer chain (Fig. 1a). Each lamin dimer bears a self-affinity, *U*_LL_, via its head and tail domains in an asymmetric way as well as an attractive domain along its rod domain (Fig. 1a). In addition, each lamin dimer can interact with the elastic shell via an interaction strength *U*_LS_ (Fig. 1b). We reason that this interaction could be weaker when the molecular coupling between the A and B meshworks is disrupted (e.g., by decreasing the expression levels of B-type lamins) or could be excessively strong (e.g., when lamin A has a mutation favoring interactions with INM, such as in progeria) [2, 8, 9, 28, 38–41].

We initialize simulations with lamin-A dimers confined to the nuclear interior, without inter-lamin or lamin–shell interactions (Fig. 1c, gray curve). In this limit, chains are broadly distributed throughout the nucleoplasm but show a mild entropic enrichment near the nuclear periphery [51], while remaining approximately homogeneous and unable to stably localize at the nuclear surface (Fig. 1d-e, gray curve). Introducing a weak interaction between lamin A dimers and the nuclear shell, comparable to thermal energy, leads to partial adsorption, with approximately seventy percent of dimers accumulating near the nuclear boundary (Fig. 1c, light blue curve). This is reflected by a sharp peripheral peak in the radial probability profile (Fig. 1d, light blue curve). Despite this enrichment, no connected network forms at the nuclear surface (Fig. 1c, light blue curve). Only when attractive interactions between lamin A dimers dominate over lamin–shell interactions do lamin A dimers assemble into a meshwork-like layer beneath the nuclear envelope (Fig. 1c-d, green curve). In contrast, introducing both lamin–lamin and lamin–shell interactions simultaneously from the outset drives bulk aggregation within the nucleoplasm (Fig. 1c, red curve and Supplementary Fig. S1). This is reflected in a reduced peripheral peak and strong accumulation of lamin chains in the nuclear interior in the radial density profiles (Fig. 1d-e, red curve). However, given that our operating lamin concentration is significantly higher than the bulk concentration, such aggregates are more likely to form near the periphery. Similar aggregation patterns both in bulk and at the periphery have been reported for certain lamin A mutants [42].

Overall, our results suggest that stable peripheral lamina formation requires a sequential assembly pathway, potentially regulated by phosphorylation–dephosphorylation cycles of lamin A and/or additional binding partners [47]. Such regulation would ensure proper segregation of lamin populations between the nucleoplasm and the nuclear periphery under physiological conditions (Supplementary Fig. S2).

### B. Competition between lamin self-association and peripheral adsorption governs lamina architecture

Having established that peripheral localization of lamin A is required for assembly of a polymer layer beneath the shell (Fig. 1), we next examined how interactions between lamin-A dimers regulate the topology of the resulting lamina meshwork (Fig. 2). Self-assembly of filamentous and branched networks from elongated proteins typically relies on attractive interaction sites that promote parallel and longitudinal associations between neighboring molecules [52–55]. Consistent with this principle, *in-vitro* studies suggest that when lamin dimers undergo a hierarchical assembly of high-aspect ratio structures (Supplementary Fig. S2), the average distance between tail domains is shorter than the full contour length of an individual dimer [18, 19, 23, 56].

To explore how such interactions influence lamina organization, we introduced an additional attractive site along the lamin dimer that mediates parallel association between neighboring dimers (see Methods). This central site interacts with both head and tail domains with affinity *U*_C_ (Fig. 1a). By varying its position relative to the terminal domains, we generated four distinct molecular architectures satisfying Δ*l*_T_ = (*n*_lamin_ *−* 1) *−* Δ*l*_H_, with 0*σ ≤* Δ*l*_H_ *≤* 3*σ* (equivalently 4*σ ≤* Δ*l*_T_ *≤* 7*σ*) (Fig. 3a). Here, Δ*l*_T_ and Δ*l*_H_ are the distances from the tail and head bead, respectively.

In the absence of the central attraction site (or when its binding strength is weak), lamin dimers assemble primarily through head-to-tail interactions, producing a cage-like meshwork with a relatively regular lattice organization (Fig. 2a, left and Supplementary Fig. S3). Introducing the central interaction site progressively enhances parallel alignment between neighboring dimers. As the site shifts toward the center of the dimer, paracrystalline bundles form and eventually reorganize into a fibrous network architecture (Fig. 2a, left to right and Supplementary Fig. S4). Increasing the strength between these sites, emulating rod interactions between dimers, *U*_C_, further promotes filament thickening and structural heterogeneity, yielding irregular fibrous meshworks as the equilibrium state (Fig. 2b).

We next investigated the role of competition between lamin-lamin interactions and adsorption to the peripheral surface (Fig. 2c-d). To do this, we assign equivalent binding strengths to all binding sites of a dimer governing the inter-lamin interactions (i.e., *U*_LL_ = *U*_C_ = *U*_HT_) (Fig. 1a). Following dimer adsorption, lamin dimers can either diffuse along the shell surface or desorb back into the bulk, depending on the relative strengths of lamin-surface and lamin-lamin interactions (Fig. 1c and Supplementary Fig. S12). Our simulations reveal that network formation requires a threshold level of intermolecular attraction, approximately *U*_LL_ *≳ U*_LS_ (Fig. 2d). Above this threshold, lamin-lamin binding favors inter-connected fibrous network structures in the presence of the central binding site or an inter-connected lattice-like network in the absence of a central binding site (Fig. 2a-c and Supplementary Fig. S3-S5).

When the lamin-lamin binding strength *U*_LL_ is weaker than surface binding *U*_LS_, dimers remain largely isolated and fail to assemble into an interconnected network (Fig. 2d, gray region and Supplementary Fig. S6). If the adsorption of lamin dimers to the shell becomes too strong, lamin dimers, kinetically trapped at their adsorption sites, substantially reduce their lateral diffusion on the surface (Supplementary Movie S1 vs. S2). Notably, strong surface binding also distorts the shell, causing a wrinkled surface that possibly further prevents lateral movement of dimers or their fibers (Fig. 2c right column and Supplementary Fig. S28 and S7). Under these conditions, network formation is suppressed despite favorable inter-lamin interactions (Fig. 2c, right panel). Thus, if effective coupling of lamin dimers to the peripheral scaffold (e.g., B meshwork) is stronger than inter-lamin coupling, the self-organization of lamins into network structures can be disrupted.

Together, these results identify the balance between inter-lamin association and surface affinity as a primary driver of fibrous structures. Fibrous network architectures emerge only when lamin dimers interact strongly enough to self-assemble, yet they should remain sufficiently mobile to reorganize on the peripheral surface.

### C. Lamin-A-free surface gaps emerge under a weak lamin-surface binding strength

The assembly of lamin A dimers into thick, periodic fibers is driven primarily by strong inter-lamin attraction arising from the competition between longitudinal and parallel modes of association (Fig. 2). As a consequence, the characteristic mesh size—defined as the size of lamin-A-free regions—depends sensitively on the balance between lamin-lamin and lamin-shell interactions (Fig. 2d and 3). Interestingly, enlarged lamin-free gaps have frequently been reported under experimental conditions in which lamin B levels are reduced or absent [10, 15, 17, 35, 45].

We hypothesized that strong binding of lamin-A dimers to the peripheral shell limits their lateral mobility after adsorption, thereby suppressing filament coarsening and promoting a more homogeneous meshwork organization. Consistent with this picture, increasing lamin-shell affinity by 2-fold produces uniformly distributed networks with smaller lamin-A-free regions (Fig. 3a-c). Under these conditions, neighboring filaments become interconnected through additional local contacts, often mediated by several lamin dimers that bridge adjacent supramolecular assemblies (Fig. 2a-b, second row). These bridging dimers persist throughout the simulations and do not disappear at long times, indicating the formation of a kinetically stable state (Supplementary Fig. S8 and S9; Table S1). Although this trend is independent of the specific mechanism of lamin-lamin binding, molecular architectures that favor parallel aggregation consistently generate larger openings within the meshwork (Fig. 3b-c and Supplementary Fig. S10-S12).

In contrast, weakening lamin-shell binding enhances the mobility of adsorbed dimers and facilitates their incorporation into growing paracrystalline fibers (Supplementary Fig. S8c and Table S2). The resulting structures are characterized by longer filamentous structures, fewer branching points, and a heterogeneous coverage of the shell surface containing large lamin-free regions (Fig. 3d and Supplementary Fig. S13). As filaments grow longer, the effective node density decreases, further increasing the characteristic spacing between neighboring fibers (Supplementary Fig. S8-S13 and Table S1, S4-S5). Consistent with these structural changes, the distribution of gap sizes broadens substantially as lamin-shell binding strength decreases while inter-lamin attraction is held fixed (Fig. 3e, purple-to-blue curves). This behavior highlights the distinct role of shell binding in regulating lamin distribution on the surface and, consequently, the large-scale organization of the lamina meshwork (Fig. 1c) [47, 57].

Together, these results suggest that, provided lamin-lamin attraction is sufficiently strong to sustain network formation, weakening lamin-A binding to the inner surface of the nucleus systematically increases filament coarsening and enlarges lamin-A free regions within the meshwork (Fig. 3d-e). Notably, this mobility-controlled mechanism is robust to variations in lamin’s bending stiffness (Supplementary Fig. S14).

### D. Simulated lamina meshworks recapitulate experimentally observed network topologies

Super-resolution (3D-SIM) imaging studies have revealed that lamin proteins organize into network-like architectures at the nuclear periphery. In particular, 3D-SIM reconstructions of lamin isoforms in mouse embryonic fibroblasts and Stochastic Optical Reconstruction Microscopy (STORM) imaging of mammalian nuclei demonstrated heterogeneous meshwork structures whose topology depends on lamin composition [12, 27]. Notably, depletion of lamin B has been reported to produce enlarged lamin-A-free gaps and surface voids within the lamina [12, 28, 35].

To quantitatively compare simulated and experimental lamina architectures, we performed a position-based network analysis and characterized each meshwork using two topological descriptors: the average edge length per face, *l*_edge_*/R*, and the number of edges per face, *n*_edge_ (Fig. 3f and Supplementary text) [58]. These metrics have previously been used to distinguish lamina meshworks in imaging experiments [12]. Larger values of *l*_edge_*/R* indicate the formation of longer continuous fibers, whereas increasing *n*_edge_ reflects the emergence of larger mesh faces and lamin-free openings (Fig. 3g) [12, 59, 60].

Using this framework, simulated lamina meshworks separate into two distinct topological classes (Fig. 3g and Supplementary Fig. S15-S16). Networks characterized by smaller values of both *l*_edge_*/R* and *n*_edge_ consist of relatively uniform mesh openings and short interconnected fibers, closely resembling experimentally observed wild-type lamin A/B architectures (Fig. 3g, green data). In contrast, larger values of both metrics correspond to heterogeneous meshworks containing long continuous fibers and large lamin-free gaps, similar to those reported following lamin-B depletion (Fig. 3g, green data). The agreement between simulations and experiments suggests that the interplay between laminlamin and lamin-shell interactions is sufficient to explain the transition from homogeneous wild-type lamina architectures to gap-rich lamin-B-deficient configurations (Supplementary Fig. S17-S18 and Table S3 vs. S4).

Consistent with this interpretation, our analysis reveals that lamin interactions jointly regulate network connectivity and mesh opening sizes (Supplementary Fig. S15-S16). Strong lamin-shell attraction favors uniformly distributed meshworks composed of shorter fibers and smaller openings, whereas weaker shell binding promotes filament coarsening, leading to long paracrystalline fibers and heterogeneous mesh sizes (Fig. 3g, and Supplementary Fig. S15-S16).

Finally, we find that these topological classes persist in simulations of larger nuclei (Supplementary Fig. S19-S20 and Table S4-S5). Although absolute values of gap size and *l*_edge_ increase with system size, distributions along *n*_edge_ remain largely unchanged (Supplementary Table S6-S7). Thus, emergent network topologies are robust to variations in the ratio between lamin dimer length and nuclear radius, *l/R*, indicating that the architectural states identified by our model represent generic features of lamin self-assembly rather than finite-size effects.

### E. Meshwork topology affects the mechanical compression response of model nuclei

Experimental studies have revealed that lamin-A meshwork is the main contributor to the mechanical response, particularly under strong deformation [10, 49, 50, 61]. Mutations of lamin A proteins that can disrupt the structure of the lamina meshwork also alter the mechanical response of the entire cell nucleus, often leading to nuclear rupture [10, 11, 15, 37, 41, 44, 62]. Further evidence indicates that mutations within the central rod domain primarily disrupt the formation of filamentous networks both *in vitro* and *in vivo*, whereas mutations in the tail domain preferentially affect the formation of paracrystalline arrays under high-salt conditions *in vitro* [1, 15, 44]. We thus apply uniaxial deformation on our isolated nuclei models to determine the dominant lamin interactions affecting the mechanical response (Fig. 4a and Supplementary Movie S3).

In order to test whether the different lamina structures in our model correspond to experiemental observations of nuclear stability, we uniaxially compressed different simulated lamina structures. To our surprise, the lamin-shell binding affinity significantly affects the compression response of the nuclei but for vanishing inter-lamin binding affinities (Fig. 4b-c). As the nucleus is squeezed in the vertical direction, strongly surface-bound lamin-A chains reinforce the shell even though they don’t form a meshwork, creating a secondary layer of adsorbed but non-connected chains, further resisting nuclear deformations (Fig. 4b, left panel, purple and red curves and Supplementary Movie S1-S2). Weakly surface-bound lamins or no-lamin cases show the weakest response to deformation as they are more mobile on the surface (Fig. 4b, left panel, gray and blue curves, and Supplementary Movie S1–S2). Notably, while inter-lamin binding has a smaller effect on the stress response than the surface-lamin binding, as the lamin-lamin binding strength is increased, the average value of the measured stress systematically but weakly increases, suggesting that a highly-stable meshwork can contribute to a strong stress response(Fig. 4c, right panel and Supplementary Fig. S21–S23).

Since various mutations in the lamin-A protein can affect peripheral binding affinity or inter-lamin interaction [1, 11, 15, 37, 62], we decided to test various meshwork topologies resulting from abnormal lamin-lamin interactions (Fig. 4d-g and Supplementary S24–S25). We perform three successive test cases in addition to our default case where each lamin can interact both head-tail and tail/head-central domain wise; We switched off the attraction between the central domain and either one or both of the terminal domains (referred to as cases *ii* and *iii* in Fig. 4d-g). We removed all head-tail binding and keep the binding between the central domain and head/tail (Fig. 4d-g, case *iv* ). Finally, we only kept the binding between the tail (or head) domain and central domain (Fig. 4d-g, case *v* ) Our simulations reveal that, consistent with our previous results (Fig. 2), omitting the attractive interactions between the central domain and either one or both terminals leads to the formation of regular, lattice-like meshworks, instead of fibrous networks (Fig. 4d-f, cases ii and iii). On the contrary, removing *U*_HT_ alone still allows for fiber formation, but these fibers are thicker, more elongated, and produce drastically large mesh openings (Fig. 4d-f, case iv). These paracrystalline arrays form nematic sheets on the surface if more lamin chains localize to the surface (Supplementary Fig. S26) [33]. Last but not least, if only a single asymmetric interaction exists between the central and one of the terminal domains, lamin dimers form surface clusters with no apparent network configuration within the meshwork (Fig. 4d-f, case v). Therefore, we conclude that head-tail interactions supplemented by at least one asymmetric interaction of the additional attractive domain along the lamin dimers are key to forming fibrous lamin supramolecular structures.

When we apply our compression protocol to the nuclei with deficient interactions, we observe that those nuclei with either meshwork or lattice-like lamina can withstand higher forces compared to the no-lamin cases (Fig. 4d-g). In these cases, the stress generated by the compression is transmitted to the lamina meshwork resisting against the lateral expansion of the nucleus (Fig. 4g). Nevertheless, when the meshwork structures lose their percolation pattern or lamin-free gaps encompass a major fraction of surface area, the resulting force response is significantly lower (Fig. 4d and g). Thus, our deformation simulations demonstrate that mutations that can disrupt lamina assembly can also decrease the mechanical integrity of the nucleus, albeit strongly surface-immobilized lamins can provide additional support against mechanical deformation.

## III. DISCUSSION

The nuclear lamina is often portrayed as a structural scaffold that supports nuclear shape and mechanics. However, accumulating experimental evidence suggests that it is better viewed as a dynamic supramolecular assembly whose organization continuously emerges from interactions among lamin proteins and their nuclear environment [25, 27, 63, 64]. Although numerous studies have characterized the molecular architecture of lamins and documented their roles in genome organization, mechanotransduction, and disease, a central question has remained unresolved: how do molecular-scale interactions between lamin dimers generate the diverse meshwork topologies observed in healthy and diseased nuclei? Here, we address this question using an explicit polymer moedel in which lamin dimers self-assemble into higher-order structures under nuclear confinement. Our results support a physical picture in which lamina architecture emerges from the interplay between adsorption to the nuclear boundary and multivalent lamin interactions. Consequently, disease-associated nuclear morphologies can be interpreted as alternative self-assembled topological states of the lamina.

A central finding of this work is that the topology of the lamina is governed by the balance between filament elongation and local network connectivity (Fig. 2). Head-to-tail interactions facilitate network connectivity but cannot by themselves generate extensive filament formation or higher-order ordering (i.e., bundling). Conversely, lateral interactions between lamins promote filamentous and paracrystalline formations (Fig. 2a-b). Varying the relative strength of these interactions generates a spectrum of architectures ranging from homogeneous meshworks of short or elongated fibrous structures and paracrystalline arrays (Fig. 2b). Notably, unlike previous continuum, network-based, or prescribed-meshwork descriptions of the nuclear lamina [49, 65– 68], our framework allows lamina topology to emerge spontaneously from the underlying interaction rules between lamins. The resulting fibrous meshwork structures closely resemble lamin networks observed in cells expressing a single lamin isoform and in several pathological conditions (Fig. 3g, red data) [1, 11, 12]. In contrast, assembly dominated by terminal interactions generates more homogeneous networks with smaller mesh openings with lattice-like structure, consistent with topologies reported for wild-type nuclei(Fig. 3g, green data) [25, 26, 69]. These observations suggest that much of the architectural diversity of nuclear lamina may arise from relatively subtle changes in the balance between terminal and lateral assembly pathways of lamin dimers. From this perspective, certain disease phenotypes, such those observed in laminopathies, arise not solely from weakened mechanical integrity but also from changes in network topology generated during lamina self-assembly.

Our result further identifies adsorption-mediated confinement as a major regulator of lamina organization. During post-mitotic nuclear assembly, lamin A accumulates at the nuclear periphery [47, 70]. Our results indicate that this localization step is not merely spatial but mechanistically important. Strong interactions with the underlying nuclear boundary can restrict lateral mobility and favor local node formation, producing uniform meshworks with relatively uniform face-size distributions (Fig. 2c). In contrast, increased surface mobility (e.g., due to the absence of lamin B ) promotes filament elongation, paracrystalline ordering, and the emergence of larger mesh openings (Fig. 3d-e and Supplementary Fig. S8). These observations are consistent with diffusion-controlled assembly mechanisms observed in other confined polymer and protein systems [52, 53, 71].

This interpretation may also provide insight into the structural differences observed between healthy and lamin-B-depleted nuclei (Fig. 3g). In the presence of a lamin-B, lamin A likely assembles underneath a relatively crowded and heterogeneous adsorption surface provided by B-type lamin meshwork, limiting lateral diffusion and promoting the formation of locally connected networks (Fig. 3g, green data). In contrast, depletion of lamin B may increase the effective mobility of lamin A at the nuclear boundary, enabling filament elongation and paracrystalline ordering(Fig. 3g, red data) [12]. Interestingly, this perspective also offers a possible explanation for observations involving progerin. Progerin retains a farnesyl group that permanently anchors it to the inner nuclear membrane [2, 9]. One might therefore expect stronger membrane attachment to suppress mobility (Supplementary Fig. S8). However, lipid membranes are intrinsically fluid structures and diffusion within the membrane plane may remain less restricted than diffusion underneath a dense lamin-B meshwork [72, 73]. Consequently, membrane anchoring alone does not necessarily imply reduced lateral mobility. Instead, the topological consequences of progerin may emerge from the combined effects of altered membrane interactions and preserved inter-lamin attractions that promote paracrystalline assembly. Although direct experimental measurements will be required to test this hypothesis, our results suggest that changes in lamin mobility may represent an under-appreciated regulator of lamina organization.

Another important implication of our calculations is the formation of lamin-free regions and mechanically vulnerable nuclear domains (Fig. 4). Lamin-free gaps frequently precede nuclear rupture, blebbing, and genome instability in both physiological and pathological contexts [10, 17, 37]. Our simulations demonstrate that weakening lamin-surface interactions can substantially increase mesh opening size and, under certain conditions, generate extensive lamin-depleted regions (Fig. 3 and 4d-g). These findings support the notion that lamin B functions as a *stabilizing adsorption substrate* for lamin A that promotes homogeneous lamin-A coverage of the nuclear surface. Loss of this stabilizing influence alters both lamin mobility and network connectivity, thereby increasing the likelihood of topological defects. Such defects are expected to act as local weak points that concentrate mechanical stress under chromatin-generated and cytoskeletal forces [15, 35, 39, 62]. Within this framework, nuclear blebs and rupture events may arise not simply from reduced nuclear stiffness but from localized structural vulnerabilities encoded within the topology of the lamina itself.

Overall, our findings support a view of the nuclear lamina as a self-organizing material whose topology encodes mechanical function. Similar to cytoskeletal networks and extracellular matrices, the lamina occupies a regime in which microscopic interaction rules determine emergent material properties. The remarkable diversity of nuclear phenotypes associated with *lmna* (lamin A) mutations may therefore reflect the sensitivity of this assembly process to relatively small perturbations in interaction strength, adsorption, or molecular mobility [15, 33, 44]. This perspective shifts attention away from individual molecular defects toward the physical principles governing network organization across scales.

Several limitations of the present model should be acknowledged. We focus on equilibrium self-assembly of a single lamin species and do not explicitly include chromatin tethering, active remodeling processes, phosphorylation-dependent turnover, or molecular differences between A- and B-type lamin networks. Furthermore, chromatin organization, membrane composition, and nuclear mechanics are all likely to influence assembly dynamics in living cells. Nevertheless, the simplified framework employed here enables isolation of the fundamental physical principles governing lamin self-organization. Future incorporation of chromatin, membrane remodeling, and active processes will enable direct investigation of how lamina topology couples to genome organization, mechanotransduction, and nuclear mechanics.

In summary, we propose that nuclear lamina architecture emerges from a competition between peripheral adsorption and multivalent lamin interactions. Such a mechanism unifies diverse experimental observations and suggests that lamina phenotypes in disease correspond to shifts between alternative self-assembled topological states of the lamina. By establishing an explicit polymer model of lamin self-assembly, our work provides a mechanistic bridge between molecular interactions and emergent nuclear architecture, offering a foundation for understanding how nanoscale perturbations propagate to cellular-scale defects in nuclear morphology, mechanics, and disease.

## VI. METHODS

### A. Molecular model of nuclear lamina

In molecular dynamics (MD) simulations, the coarse-grained mesoscopic nucleus model consists of three components: i) short semiflexible chains representing lamin A dimers (Fig. 1a); ii) long flexible homopolymers emulating the osmotic pressure caused by chromatin (Fig. 1b); and iii) a bead-spring elastic shell providing a fluctuating surface for the lamina assembly (Fig. 1b).

Since the smallest assembly unit of lamin fibers is a coiled dimer of length *l ≈* 50 nm, we model lamin dimers as a monodisperse solution of coarse-grained semiflexible chains in implicit solvent (Fig. 1a) [18, 74]. Each dimer is composed of *n*_lamin_ = 8 monomers, and the ratio of persistence length to contour length is fixed to *l*_p_*/l* = 2.5, ensuring semiflexibility. To model their head-to-tail assembly, each lamin is composed of attractive head and tail domains, and a central hydrophobic region (Fig. 1a, *U*_HT_). We also introduce an additional attractive domain within the central hydrophobic region, responsible for the lateral alignment of lamins when they form higher-order structures such as protofilaments or thicker fibrous structures in disease (Fig. 1a, *U*_C_) [23, 25, 44]. Together, the head-tail and lateral interactions between the head, tail, and additional attractive domains constitute inter-lamin interactions, *U*_LL_ (Fig. 1a).

*In vivo* studies indicate that the copy number of lamin proteins within the nucleus is approximately 10^6^ with a bulk concentration of around 100 nM *−* 1*µ*M [61, 75, 76]. Assuming a nuclear diameter of *d ≈* 10 *µ*m, such concentrations lead to a nucleoplasmic-lamin volume fraction less than *≈* 1%. However, lamin concentration near the nuclear lamina can be much higher, exceeding several tens of *µ*M. Given the approximate nature of stoichiometric metrics for lamins, we choose a lamin volume fraction of *ϕ ≈* 24% with *N*_lamin_ = 1250 lamin chains for our smallest nuclear shell with radius *R* = 34*σ*, where 1*σ* is the unit monomer size in simulations. This choice also accounts for lamina thickening and lamin overexpression in disease, as well as nucleoplasmic lamins distributed throughout the bulk of the nucleus [8, 9, 11, 77]. In other words, upon assembly, the lamina occupies 40-90% of the shell surface (Supplementary Fig. S13, *U*_LL_ *>* 6.0*k*_B_*T* ), encompassing the experimentally observed coverage of *≈* 50*±*10% of the nuclear surface by lamin filaments [25]. Notably, this concentration range also corresponds to *c ≈* 2.0*c*^*\**^, where *c*^*\**^ is the bulk overlap concentration. The overlap concentration, at which lamin chains begin to interact, for *n*_lamin_ = 8 monomers is *c*^*\**^ = 0.03*σ*^*−*3^.

To model interphase chromosomes, each nucleus contains *n*_polymer_ = 4 flexible chains at the interior, occupying a volume fraction of *≈* 10% [66, 78, 79] (Fig. 1b and Supplementary Fig. S27b). These homopolymers prevent the inward collapse of the shell and represent the osmotic pressure generated by chromatin within the nucleus. Chromatin density stays uniform during data acquisition, after initial polymer relaxation steps [66, 80].

At the beginning of a simulation at *t* = 0, both flexible polymers and semiflexible dimers are placed within the elastic shell (Fig. 1b and Supplementary Fig. S27a). The elastic shell is composed of at least *N*_shell_ = 22500 beads. Each bead forms an average of *⟨n*_bond_*⟩* = 5.5 bonds, with appropriate constraints to prevent bead overlap, steric clashes, or large openings within the shell, as described in our previous work [65, 66, 80]. Since the average lamin filament size, *l*, and nuclear diameter, *R* are *l/R ≈* 400 nm*/*5 *µ*m *≈* 0.04 - 0.1) [25], we also performed simulations with larger nuclear shells and observed qualitatively similar results (Supplementary Fig. S2).

### B. MD simulation setup

Initially, lamin chains and chromatin homopolymers are placed randomly within the elastic nuclear confinement (Fig. 1c and Supplementary Fig. S27a). An energy minimization process shrinks the shell by approximately 15% (Supplementary Fig. S27c). This closes large openings in the shell, preventing the escape of interior contents (i.e., lamin chains and flexible polymers) (Supplementary Fig. S27c). This is followed by a relaxation step for 500*τ* with a larger, default time step of Δ*t* = 0.005*τ* . The time unit is defined as [*τ* ] = (*mσ*^2^*/ϵ*)^1*/*2^, where *m* = 1 is the reduced mass of each monomer, and *ϵ* is the energy unit in our simulations. Relaxation is purely repulsive, allowing both flexible polymers and semiflexible lamin chains to mix isotropically within the elastic nucleus (Fig. 1c, gray curve). Upon relaxation, attractive lamin-shell interactions are introduced to localize lamin polymers to the shell boundary for 5 *×* 10^3^*τ* with default Δ*t* (Fig. 1c, light blue curve). This ensures that *≈* 70% of all chains localize to the interior of the elastic shell (Fig. 1c, light blue curve). Finally, the data production step involves introducing inter-lamin interactions for an additional 1.25 *×* 10^4^*τ* to induce lamina meshwork formation (Fig. 1c, green curve).

All bonded interactions of the elastic shell, chromatin homopolymers, and lamin chains are modeled by a Finitely Extensible Non-linear Elastic (FENE) potential

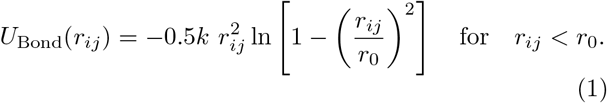

where *k* is the spring constant, and *r*_0_ = 1.5*σ* is the maximum bond length [74, 81]. We use bond energy *k* = 30.0*k*_B_*T/σ*^2^ for interior homopolymers and lamin dimers. To ensure deformability, shell beads have a lower bond stiffness of *k* = 5.0*k*_B_*T/σ*^2^ [80]. *k*_B_ is the Boltzmann constant and *T* is the reduced absolute temperature, unless otherwise stated.

A harmonic bending potential modulates the bending rigidity of lamins and homopolymers

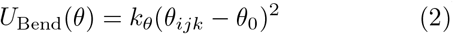

where *θ*_*ijk*_ is the angle between two bonds, *θ*_0_ = *π* is the reference angle and *k*_*θ*_ defines the bending stiffness. We set *k*_*θ*_ = *k*_B_*T/*rad^2^ for flexible polymers and *k*_*θ*_ = 20.0*k*_B_*T/*rad^2^ for lamin chains. This yields persistence lengths of *l*_p_ = 1*σ* for flexible polymers and *l*_p_ = 19.4*σ* for lamin dimers, where *l*_p_ *≈ k*_*θ*_ *l*_b_*/k*_B_*T* [74]. *l*_b_ is the average bond length, *l*_b_ *≈* 0.97*σ* [74].

All non-bonded, steric interactions within the elastic nucleus are described by a shifted and truncated Lennard-Jones (LJ) potential

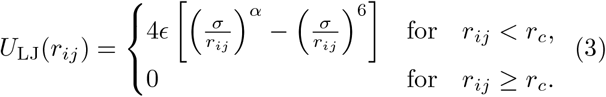

where *r*_*ij*_ is the distance between two interacting beads *i* and *j, ϵ* is the interaction strength, *σ* is the bead diameter and *α* specifies the exponent of the repulsive term. The default value of interaction strength in Eq. 3 is *ϵ* = 1*k*_B_*T* . Lamin-shell and inter-lamin binding affinities are modeled by varying the strength of the LJ potential, *ϵ*, and we report these strengths as binding interactions throughout this work (Fig. 1a). Since the physical nature of how lamin supramolecular structures form is largely elusive, we probe lamin-shell binding in the range 0.75*k*_B_*T ≤ U*_LS_ *≤* 5.0*k*_B_*T* . This range spans weak interactions, where lamin localization is largely governed by thermal fluctuations, to stronger attractions that enhance localization. Close to the upper limit (i.e., *U*_LS_ *≥* 2.5*k*_B_*T* ), lamin localization strength can distort the shape of the elastic shell (Supplementary Fig. S28). Inter-lamin attractions are varied such that 2.5*k*_B_*T ≤ U*_LL_ *≤* 25.0*k*_B_*T* to encourage network-like assembly of the lamina. These attraction ranges are consistent with previous models with similar coarse-graining levels [33, 80, 82, 83]. For all attractive binding interactions, we use a cutoff *r*_c_ = 2.5*σ* and an exponent *α* = 12. This enforces a sharper and more rigid interaction potential [80, 84–86]. In contrast, the cutoff distance for the rest of monomer-monomer interactions is *r*_*c*_ = 2^1*/*6^*σ* with an exponent *α* = 9 for the chromatin polymer or 12 for the shell and lamin beads. These parameters provide repulsive steric interactions between the chromatin polymer, shell, and lamin beads.

MD simulations are performed in the canonical ensemble (i.e., constant volume *V*, particle number *N*, and temperature *T* ), using a Langevin thermostat. All simulations are performed with the LAMMPS MD package [87, 88]. Lamina meshwork configurations are visualized using OVITO [89] and characterized using various Python libraries.

### C. Characterizing lamina meshwork phases

Lamina meshwork topologies obtained in our simulations are characterized by a position-based network methodology (Supplementary Fig. S29-S30). This is analogous to computational image analysis previously used to reconstruct 3D structured illumination microscopy (3D-SIM) of the lamina [12, 59, 60] (Supplementary text).

To do so, the lamina meshwork configuration is first converted into a node-edge graph that preserves its spatial topology [90, 91] (Supplementary Fig. S29a). This is followed by a clustering algorithm, where all neighboring beads within a cut-off radius, *r*_*merge*_ *<* 5.5*σ* (i.e., *≈* 0.7 *· n*_lamin_) are clustered to form a single node (Supplementary Fig. S3a and S4a). In other words, *r*_*merge*_ is the threshold that yields a connected lamina network with 150 *≤* node count *≤* 850 (Supplementary Fig. S31). A merge distance above the threshold, *r*_merge_ *>* 5.5*σ*, results in non-network configurations, where either the meshwork fails to achieve connectivity or leads to over-connectivity (Supplementary Fig. S32-S33). With this algorithm, we obtain simplified 2D reconstructions of the lamina, retaining only its mesoscale connectivity (Supplementary Fig. S30c and S4b) [12].

Upon network graph reconstruction, we extract structural properties such as average node count, number of edges per face, *l*_edge_, and edge length per face, *n*_edge_, to quantify the resulting meshwork [58] (Fig. 2d, 3f-g and Supplementary Fig. S30d). These metrics have been previously used to quantify lamina meshworks visualized by three-dimensional structured illumination microscopy (3D-SIM) [12] (Supplementary text).

While a 2D reconstruction is sufficient to obtain cluster metrics mentioned above, quantifying the characteristic face size for each meshwork would require 3D analysis over the elastic nucleus (Supplementary Fig. S34). Hence, we employ a geometric filling procedure that directly identifies lamin-free regions, or meshwork faces, on the shell surface (Fig. 3a).

The lamina meshwork face size distribution is obtained by projecting the meshwork onto a best-fit sphere and performing random sequential adsorption (RSA) process [92] (Supplementary text). Each point on the surface is sampled to insert a new bead, such that it does not overlap with any existing lamin beads [93]. Neighboring newly-added beads are then grouped using a graph-based clustering algorithm [94]. The face area, *A*_face,i_, is then calculated as

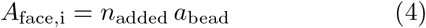

where *n*_added_ is the number of beads per face, and *a*_bead_ is the area of each bead. Average face area, *A*_face_, is calculated as the weighted average of the face size distribution for each simulated meshwork (Fig. 3).

To further characterize lamin surface coverage in the meshwork, we also utilize the solvent accessible surface area (SASA) metric to calculate lamin-covered area (LCA) [95] (Fig. 3a, purple region and Supplementary Fig. S35). This metric yields trends that are the inverse of *A*_face_, validating our geometry-filling methodology for computing mesh opening sizes, as described above.

## Supporting information

Supplementary Information for "Multivalent lamin binding controls the meshwork structure in a self-assembled model of the nuclear lamina"

## DATA AVAILABILITY

All codes, structure files, and input parameters are available online at https://github.com/erbaslab.

## ACKNOWLEDGMENTS

We acknowledge Edward J. Banigan, Andrew D. Stephens, Abigail Buchwalter, Alexander von Appen, and Michele Nava for fruitful discussions. This research was supported by TUBITAK, the Scientific and Technological Research Council of Turkey, under TRANSCAN 3 (SCIE-PANC) [Grant No. 124N935]. The simulations were partially performed at TUBITAK ULAKBIM High Performance and Grid Computing Center (TRUBA).

