## Supplementary Information for "Multivalent lamin binding controls the meshwork structure in a self-assembled model of the nuclear lamina" for "Multivalent lamin binding controls the meshwork structure in a self-assembled model of the nuclear lamina"

### 1 Concentration of semiflexible chains, $c$

In our coarse-grained nuclear lamina model, the concentration of lamin A dimers within the bonded elastic shell is fixed to $c \approx 0.06\sigma^{-3}$, which corresponds to $N_{\text{lamin}} = 1250$ chains within the bonded elastic shell. $c$ is calculated as follows

$$c = \frac{N_{\text{lamin}} \cdot n_{\text{lamin}}}{V} = \frac{1250 \cdot 8}{\frac{4}{3}\pi(34\sigma)^3} \approx 0.06\sigma^{-3} \tag{1}$$

where $n_{\text{lamin}} = 8$ is the number of monomers per lamin dimer, and $V$ is elastic shell volume.

We also denote $c$ in terms of bulk overlap concentration, $c^*$, where lamin chains begin to interact with one another. For $n_{\text{lamin}} = 8$, $c^*$ is as follows

$$c^* = \frac{n_{\text{lamin}}}{v} = \frac{8}{\frac{4}{3}\pi(4\sigma)^3} = 0.02984155183 \approx 0.03\sigma^{-3} \tag{2}$$

where $v \approx l\sigma^3$ is the pervaded volume of each chain.

Absolute concentration of lamin chains, $c$, can then be denoted as

$$c = \frac{c}{c^*} \approx \frac{0.06}{0.03} \approx 2.0c^* \tag{3}$$

This concentration sufficiently covers the elastic shell surface to form network and occupies 24% of the elastic shell volume to account for lamin overexpression or accumulation in disease [1] (see main text).

### 2 Meshwork analysis metrics

To quantitatively characterize the lamina meshwork topologies observed in our simulation, we use several metrics such as geometric metrics, characteristic mesh opening size, $A_{\text{face}}$, and lamin-covered area, LCA. These are described in detail below.

#### 2.1 Meshwork reconstruction and geometric merging

We develop a geometric reconstruction-based analogue for computational image analysis of 3D structured illumination microscopy (3D-SIM) images [2]. Nevertheless, such analysis can be applied to any particle-based meshwork trajectory. Our reconstruction is an averaged configuration over the trajectory from the last half of the lamina assembly step (Fig. 1c). It follows the following procedure:

1. **Mapping lamina topology to geometric graph:** The coordinates of each bead that constitutes the lamina are mapped onto a geometric graph by treating the bead positions as a point cloud in 3D Euclidean space [3, 4] (Fig. S4a).

2. **Preserving spatial connectivity of initial lamina network:** Edges are defined between the points such that the spatial proximity of the initial network structure is preserved (Fig. S4a). Edges between two beads are defined $i$ and $j$ such that

$$\| r_i - r_j \| \leq r_{\text{cut}}$$

where a cut-off distance, $r_{\text{cut}} = 1\sigma$, corresponding to the characteristic bead size. This is chosen to be slightly larger than the average bond length, $l_{\text{b}} \approx 0.97\sigma$. This is to ensure that directly bonded neighbors and near-contact interactions of each are accurately captured.

3. **Network clustering:** The node-edge geometric network, constructed in previous steps, is then used to identify local neighborhoods within a merge radius, $r_{\text{merge}}$, such that

$$r_{\text{merge}} = \beta\, r_{\text{cut}}$$

where $\beta$ is chosen as the smallest coefficient that yields a fully connected network. Neighboring beads within $r_{\text{merge}}$ are then replaced by a single coarse-grained node located at the center of mass of the beads in that local neighborhood. In our simulations, $\beta_{\text{max}} = 5.5\sigma \approx 0.7 \cdot n_{\text{lamin}}$, beyond which we observe non-network lamina meshworks (see main text).

4. After each merging step, edges are redefined between the newly formed nodes based on the original adjacency. The process is repeated iteratively while $r_{\text{merge}} \leq \beta_{\text{max}}$, eventually yielding a fully connected network, or non-network favoring topologies (Fig. 2c). Network connectivity is preserved by reassigning all preexisting edges to the corresponding geometric clusters, ensuring that the large-scale topology of the meshwork remains intact (Fig. S4c).

Constructing a geometric network allows us to corroborate experimentally observed healthy, knockout, and diseased lamina meshworks and determine their topological features and interactions [5, 6, 7] (Fig. S4d). Hence, we choose experimentally described quantities that characterize network connectivity, and surface accessibility. In particular, we employ average node count, average edge length per face, $l_{\text{edge}}$, and number of edges per face, $n_{\text{edge}}$.

##### 2.1.1 Average node count

Average node count is a time-averaged quantity representing the global connectivity of the meshwork. In our simulations, average node count spans the range $200 \leq$ Average node count $\leq 850$. Node counts outside this range emerge only for weakly connected or disconnected networks, where $r_{\text{merge}} > 5.5\sigma$.

##### 2.1.2 Number of edges per face, $n_{\text{edge}}$

Number of edges per face, $n_{\text{edge}}$, is an indirect measure of face size and reflects changes in network connectivity and face shape in particular. It is computed as the total edge count, $n_{\text{e}}$, within our 3D network over the number of faces, $n_{\text{f}}$ obtained from its 2D planar representation (see Fig. S4 and S7).

$$n_{\text{edge}} = \frac{n_{\text{e}}}{2 \cdot n_{\text{f}}} \tag{4}$$

##### 2.1.3 Average edge length per face, $l_{\text{edge}}/R$

Average edge length per face, $l_{\text{edge}}/R$, is defined as the mean Euclidean length of all edges bounding a given face. Hence, it indicates the formation of long continuous fibers as it increases ($l_{\text{edge}}/R > 0.34$), and is often low for uniform meshworks ($l_{\text{edge}}/R < 0.34$). Note that $l_{\text{edge}}/R$ is a dimensionless quantity and is normalized by the radius of gyration of the bonded elastic shell ($R_{\text{g}}$. This is to exclude size effects, similar to experiments, where the measure is normalized by nuclear size [2].

$$l_{\text{edge}}/R = \frac{l_{\text{total}}}{2 \cdot n_{\text{f}}} \cdot \frac{1}{R_{\text{g}}} \tag{5}$$

where $l_{\text{total}}$ is the total Euclidean edge length constituting the geometric network.

*Other network features, such as node assortativity, centrality, average shortest path between nodes, etc., can also be obtained from our network analysis algorithm.

#### 2.2 Meshwork face size, $A_{\text{face,i}}$

It is important to note that while the meshwork reconstruction methodology is suitable for extracting clustering-based metrics such as average node count, $n_{\text{edge}}$, and $l_{\text{edge}}$, quantifying face size distribution, $A_{\text{face,i}}$, would require projecting the 3D geometric graph to a hemisphere in $xy$ plane (Fig. S6). Therefore, we employ the random sequential adsorption (RSA) algorithm to geometrically fill lamin-free regions.

1. **Meshwork projection onto perfect sphere:** The simulated lamina meshwork is first projected onto a best-fit spherical surface. The radius and center of mass of the sphere is determined by minimizing the mean squared deviation between the bead positions and the sphere.

2. **Surface sampling and geometric filling:** Once all meshwork beads are projected onto the surface, we employ the random sequential adsorption (RSA) process. Every point on the surface is sampled to insert a new bead, while enforcing a minimum separation of $d_{\text{bead}} = 1\sigma$ from existing beads. In other words, $d_{\text{bead}} < 1\sigma$ indicates the presence of a lamin bead. With this constraint, additional beads only fill locations unoccupied by lamins (i.e., meshwork faces).

3. **Calculating face area:** Neighboring newly-added beads are then grouped using a graph-based clustering algorithm [8]. The number of beads per cluster, $n_{\text{added}}$ is used to calculate the face size, $A_{\text{face,i}}$, as

$$A_{\text{face,i}} = n_{\text{added}}\, a_{\text{bead}} = n_{\text{added}}\, \pi \left(\frac{d_{\text{bead}}}{2}\right)^2 \tag{6}$$

   where $a_{\text{bead}}$ is the area of each bead. Average face size, $A_{\text{face}}$, is then calculated as the weighted mean of the face size distribution.

##### 2.3 Lamin-covered area (LCA)

Lamin-covered area (LCA) is another indirect measure of the lamina meshwork opening size, derived from solvent-accessible surface area (SASA), and hence an inverse to mesh opening size. In other words, a meshwork with large opening sizes would result in low LCA and vice versa. LCA is computed via a sphere with a radius of $0.5\sigma$ probing the meshwork surface, yielding an effective surface area covered by lamin dimers. Regions not covered by lamins fibers mark the solvent-excluded surface (gray). This metric yields trends inverse of $A_{\text{face}}$, serving as a validation to our geometry filling methodology to compute mesh opening sizes, described above. We also obtained LCA normalized by the radius of gyration of the nuclear shell to obtain comparable values across various system sizes.

$$\text{LCA}_{\text{norm}} = \frac{\text{LCA}}{R_g^2} \tag{7}$$

#### 3 Normalized asphericity, $A_{\text{norm}}$

As discussed in the main text, high lamin-shell attraction ($U_{\text{LS}} = 5.0 k_B T$) leads to distorted elastic shell shapes. These shells fail to project onto a uniform sphere, and hence, lead to isotropic topologies, which we mark as non-network cases in Fig. 3b (see main text). To quantitatively confirm this, we calculate the normalized asphericity of the bonded elastic shell (see Fig. S3 and S15).

$A_{\text{norm}}$, is a normalized version of asphericity, $A$, a scalar metric obtained from the elastic shell's gyration tensor to quantify its deviation from a perfect sphere [9]. $A$ is normalized as follows

$$A_{\text{norm}} = \frac{A}{R_g^2} = \frac{\lambda_1 - \frac{1}{2}(\lambda_2 + \lambda_3)}{R_g^2} \tag{8}$$

where $\lambda_1$, $\lambda_2$, and $\lambda_3$ are eigenvalues of the gyration tensor of the elastic shell, $S$, and $R_g$ is the radius of gyration of the shell (see main text). Dividing by $R_g^2$ makes $A_{\text{norm}}$ dimensionless, meaning that it eliminates the effect of shell size on asphericity.

### 4 Uniaxial compression simulations

To apply uniaxial deformation in simulations, we compress the elastic nuclei after the network formation step along the $z$-axis. Two static, repulsive bead plates were initially placed at $z$ = $\pm 70\sigma$ (i.e., initial length of $L_{z,i} = 140\sigma$). They were moved to $z = \pm 15\sigma$ (i.e., final length of $L_{z,f}$ = $30\sigma$) over $10^5$ timesteps, which corresponds to $100\tau$, with a timestep of $\Delta t = 0.001\tau$. This is followed by stress relaxation, where the parallel plates are kept at $L_{z,f} = 30\sigma$ for an additional $5 \times 10^4$ timesteps ($50\tau$). A sample stress-time curve is shown in the main text, Fig. 4a. The strain, $\varepsilon_z$, can be calculated as

$$\varepsilon_z = \frac{L_{z,i} - L_{z,f}}{L_{z,i}} \tag{9}$$

which corresponds to a final compressive strain of approximately ($\varepsilon_z \approx 0.79$) (79% compression).

### 5 Supplementary figures and tables

Table S1: Average node count does not alter significantly upon doubling simulation time, as shown visually in Fig. S17a.

| $U_{LS}(k_B T)$ | $t = 1.25 \times 10^4 \tau$ | $t = 2.5 \times 10^4 \tau$ |
|---|---|---|
| 0.75 | 485 | 547 |
| 1.5 | 740 | 785 |

Table S2: Characteristic relaxation times, $\tau$, and effective diffusion coefficients, $D$, obtained from fitting the normalized positional autocorrelation function shown in Fig. S17b.

| $U_{LS}$ ($k_B T$) | $\tau$ (timesteps) | $D$ ($\sigma^2$/timestep) |
|---|---|---|
| 0.75 | $(6.15 \pm 0.01) \times 10^5$ | $9.40 \times 10^{-4}$ |
| 1.50 | $(2.16 \pm 0.08) \times 10^6$ | $2.68 \times 10^{-4}$ |

Table S3: Face area, $A_{face}$ obtained from 3D-SIM images shown in Fig. S26-S27 via ImageJ.

| | wt A | lmnb$^{-/-}$ |
|---|---|---|
| Average face area, $A_{face}$ ($\mu m^2$) | 0.303 | 1.154 |
| Standard deviation | 0.1470 | 0.5279 |
| Surface area ($\mu m^2$) | 331.346 | 265.170 |
| Avg. $A_{face}/A_{surf}$ | 0.000944 | 0.00436 |
| Max. $A_{face}/A_{surf}$ | 0.00136 | 0.00772 |
| Min. $A_{face}/A_{surf}$ | 0.000470 | 0.002361 |

Table S4: Average normalized face area, $A_{\text{face}}/A_{\text{surf}}$, for different shell radii and lamin interaction strengths. We observe an order of magnitude difference as $A_{\text{face}}$, which is qualitatively consistent with experimental values, shown in Table S2.

**$A_{\text{face}}/A_{\text{surf}}$ for default system**

| $R = 34\sigma$ | | |
|---|---|---|
| $U_{\text{LS}}$ | $U_{\text{LL}} = 10.0k_{\text{B}}T$ | $U_{\text{LL}} = 25.0k_{\text{B}}T$ |
| $0.75k_{\text{B}}T$ | 0.029123 | 0.029753 |
| $1.50k_{\text{B}}T$ | 0.002927 | 0.009810 |
| $2.50k_{\text{B}}T$ | 0.001514 | 0.003220 |

**$A_{\text{face}}/A_{\text{surf}}$ for larger system size**

| $R = 68\sigma$ | | |
|---|---|---|
| $U_{\text{LS}}$ | $U_{\text{LL}} = 10.0k_{\text{B}}T$ | $U_{\text{LL}} = 25.0k_{\text{B}}T$ |
| $0.75k_{\text{B}}T$ | 0.020951 | 0.010689 |
| $1.50k_{\text{B}}T$ | 0.000976 | 0.003450 |
| $2.50k_{\text{B}}T$ | 0.000216 | 0.000910 |

Table S5: Comparable normalized LCA values for elastic nuclei with radii $R = 34\sigma$ and $68\sigma$ with a constant surface concentration.

**Normalized LCA for default system (Fig. 2c)**

| $R = 34\sigma$ | | |
|---|---|---|
| $U_{\text{LS}}$ $(k_{\text{B}}T)$ | $U_{\text{LL}} = 10.0k_{\text{B}}T$ | $U_{\text{LL}} = 25.0k_{\text{B}}T$ |
| 0.75 | 0.5683 | 0.5683 |
| 1.50 | 0.5683 | 0.7305 |

**Normalized LCA for larger system size (Fig. S28)**

| $R = 68\sigma$ | | |
|---|---|---|
| $U_{\text{LS}}$ $(k_{\text{B}}T)$ | $U_{\text{LL}} = 10.0k_{\text{B}}T$ | $U_{\text{LL}} = 25.0k_{\text{B}}T$ |
| 0.75 | 0.5295 | 0.6224 |
| 1.50 | 0.9602 | 0.7614 |

Table S6: Comparison of metrics for healthy lamina in Fig. 3g (green data) with larger system for $U_{\text{LS}} = 1.5k_{\text{B}}T$ $U_{\text{LL}} = 10k_{\text{B}}T$.

| | $R = 34\sigma$ | $R = 68\sigma$ |
|---|---|---|
| Number of nodes | 459 | 1343 |
| Nodes/$R_g^2$ | 13.50 | 19.75 |
| Number of edges/face, $n_{\text{edge}}$ | 4.460 | 4.491 |
| Mean edge length/face, $l_{\text{edge}}$ | 9.600 | 11.617 |
| Normalized mean edge length/face, $l_{\text{edge}}/R$ | 0.2823 | 0.1708 |

Table S7: Comparison of metrics for lamin B lacking lamina in Fig. 3g (red data) with larger system for $U_{\text{LS}} = 0.75k_{\text{B}}T$ $U_{\text{LL}} = 10k_{\text{B}}T$.

| | $R = 34\sigma$ | $R = 68\sigma$ |
|---|---|---|
| Number of nodes | 400 | 819 |
| Nodes/$R_g^2$ | 11.76 | 12.04 |
| Number of edges/face, $n_{\text{edge}}$ | 6.610 | 6.613 |
| Mean edge length/face, $l_{\text{edge}}$ | 13.137 | 16.634 |
| Normalized mean edge length/face, $l_{\text{edge}}/R$ | 0.3860 | 0.2446 |

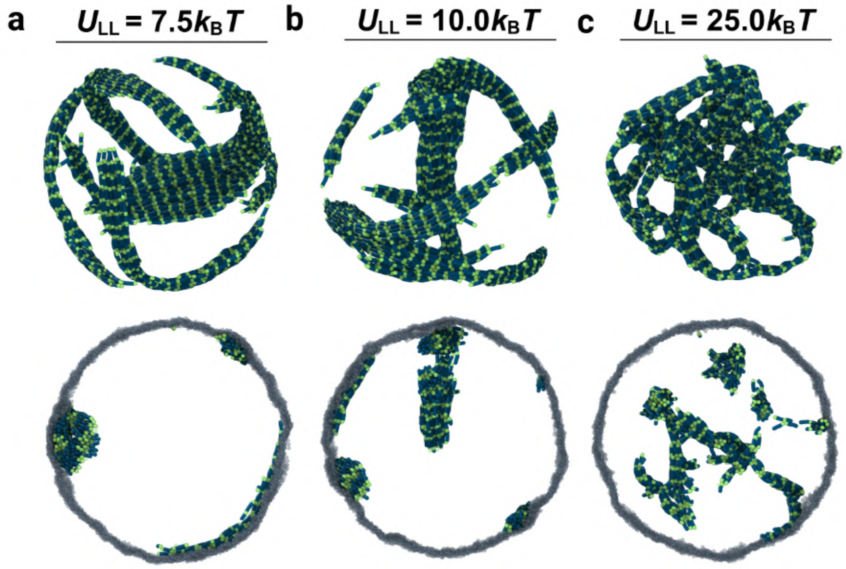


Figure S1: Introducing both inter-lamin and lamin-shell binding affinities simultaneously leads to bulk aggregates when inter-lamin affinity is sufficient to form networks, $U_{\mathrm{LL}} > 6.0 k_{\mathrm{B}} T$. Lamin-shell binding is fixed to weak, $U_{\mathrm{LS}} = 0.75 k_{\mathrm{B}} T$. Top panels show lamin chains only, while the bottom panels show a $5\sigma$ from the center of the elastic nuclear shell, showing bulk aggregates.

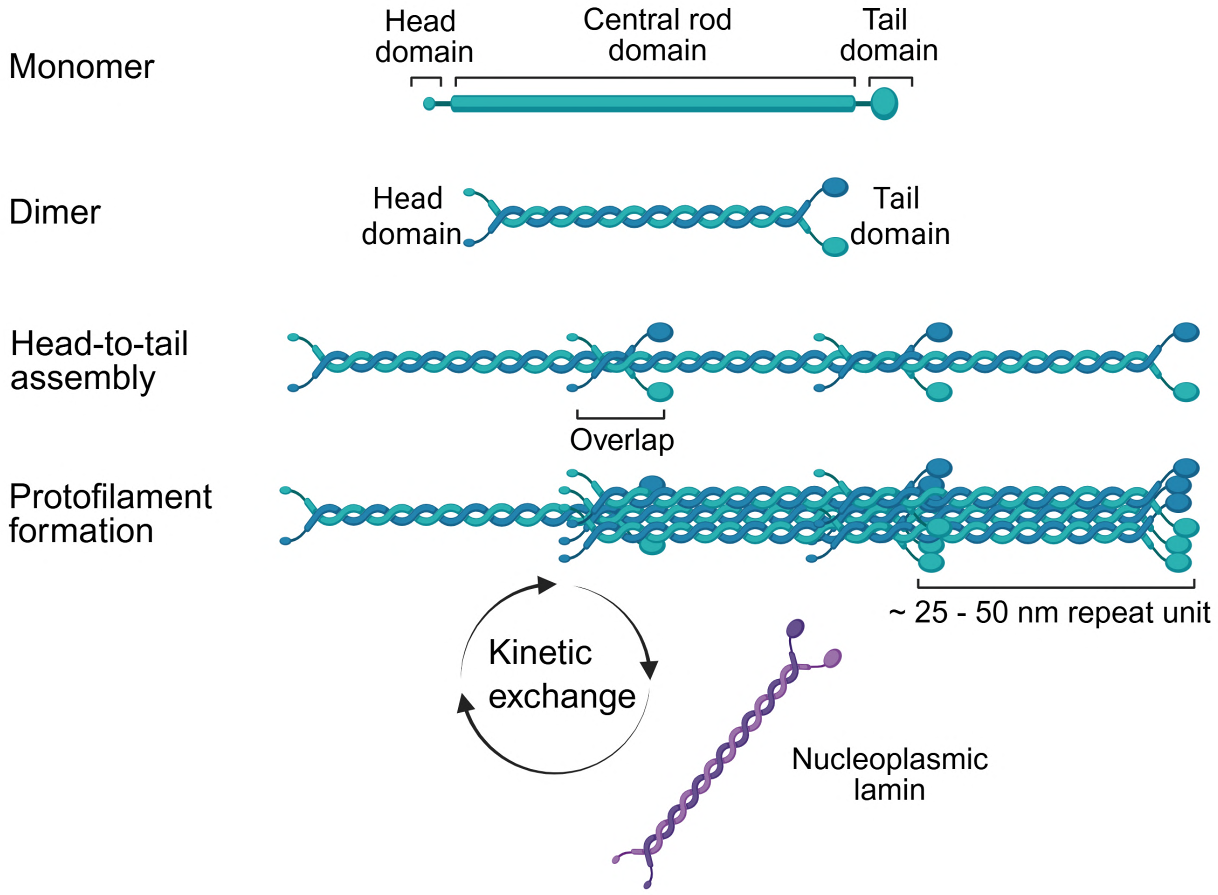


Figure S2: Schematic representation of various levels of structural assembly of lamin monomers, and how they undergo kinetic exchange with their nucleoplasmic counterparts.

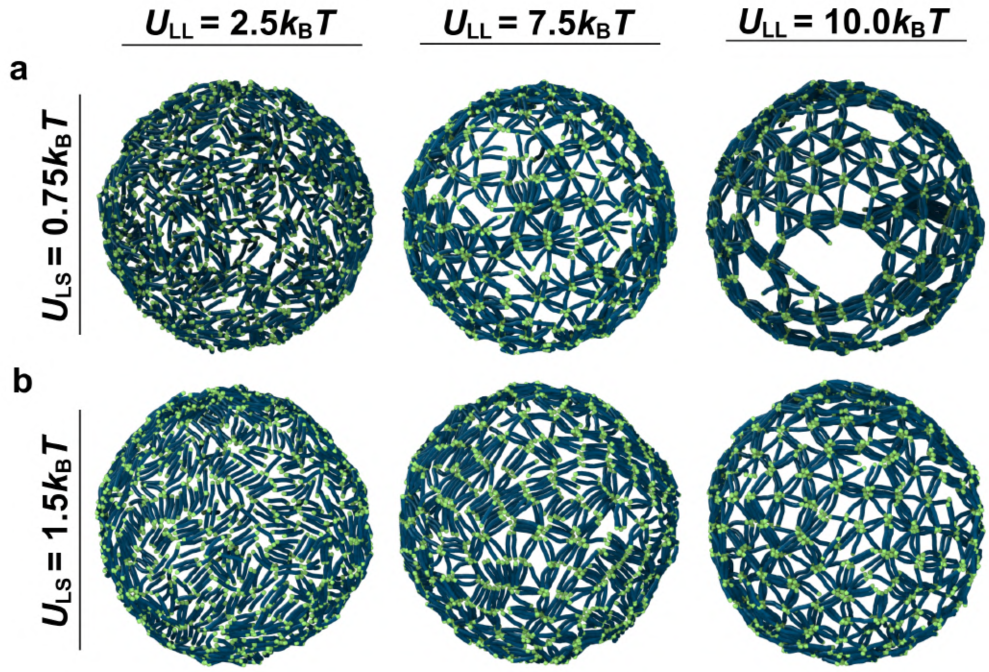


Figure S3: The absence of the additional attraction site leads to lattice-like networks instead of paracrystalline array formation (a) As expected, low lamin-shell attraction ($U_{\mathrm{LS}} = 0.75k_{\mathrm{B}}T$) leads to enlarged mesh size opening as inter-lamin attraction increases ($U_{\mathrm{LL}} = 10.0k_{\mathrm{B}}T$). (b) $U_{\mathrm{LS}} = 1.5k_{\mathrm{B}}T$ leads to uniform surface coverage.

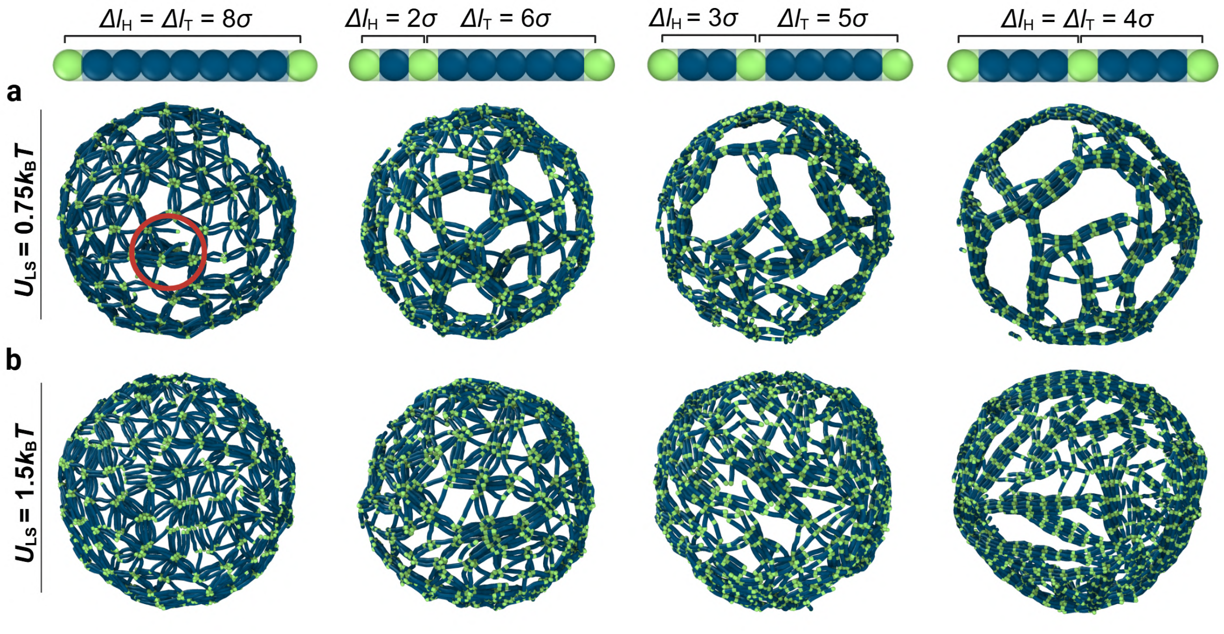


Figure S4: Lamin dimer with $n_{\mathrm{lamin}} = 9\sigma$ (symmetric $\Delta l_{\mathrm{H}}$) leads to similar meshwork phases when an additional attraction site, $C$, is at or close to the center (last two panels). Inter-lamin attraction is fixed to $U_{\mathrm{LL}} = 10.0k_{\mathrm{B}}T$. (a) Low lamin-shell attraction, $U_{\mathrm{LS}} = 0.75k_{\mathrm{B}}T$ yields paracrystalline arrays with large mesh openings, while (b) $U_{\mathrm{LS}} = 1.5k_{\mathrm{B}}T$ leads to fibrous meshwork.

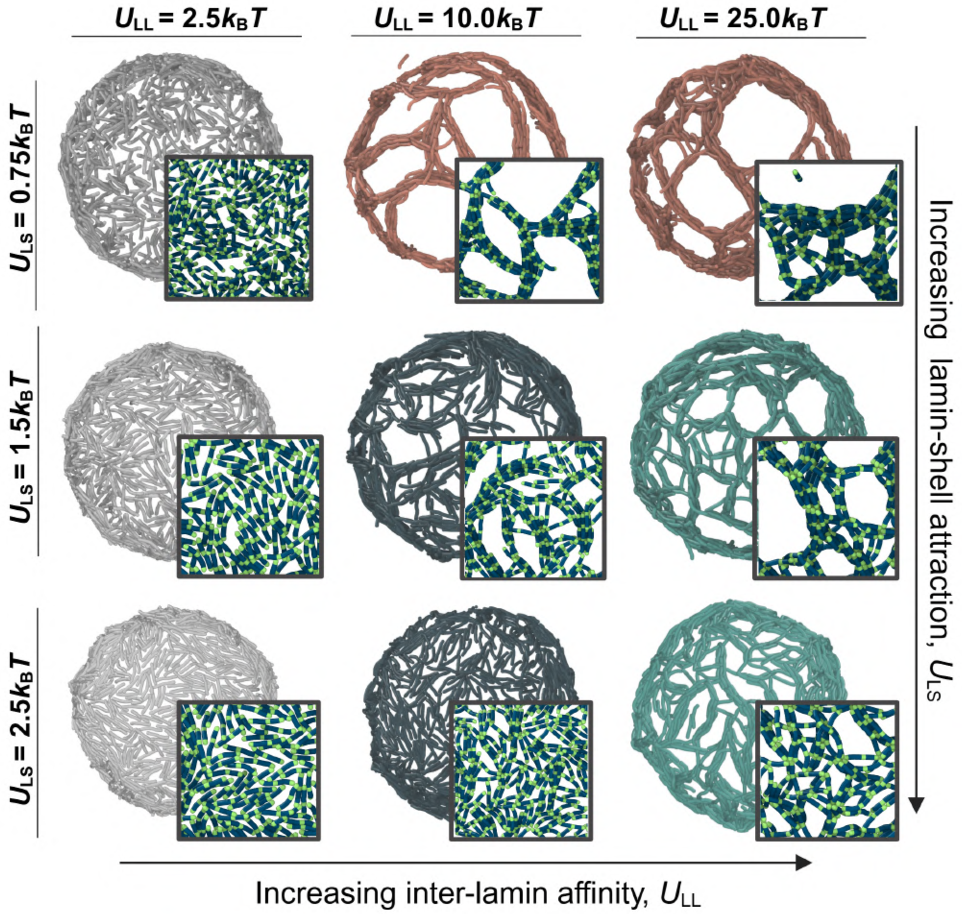


Figure S5: Interplay of lamin interactions leads to distinct meshwork phases. Color codes represent various network configurations. Gray: non-network cases, orange: paracrystalline arrays, green: networks with thin fibers, blue: largely isoptropic arrangements characterized by branched fibers.

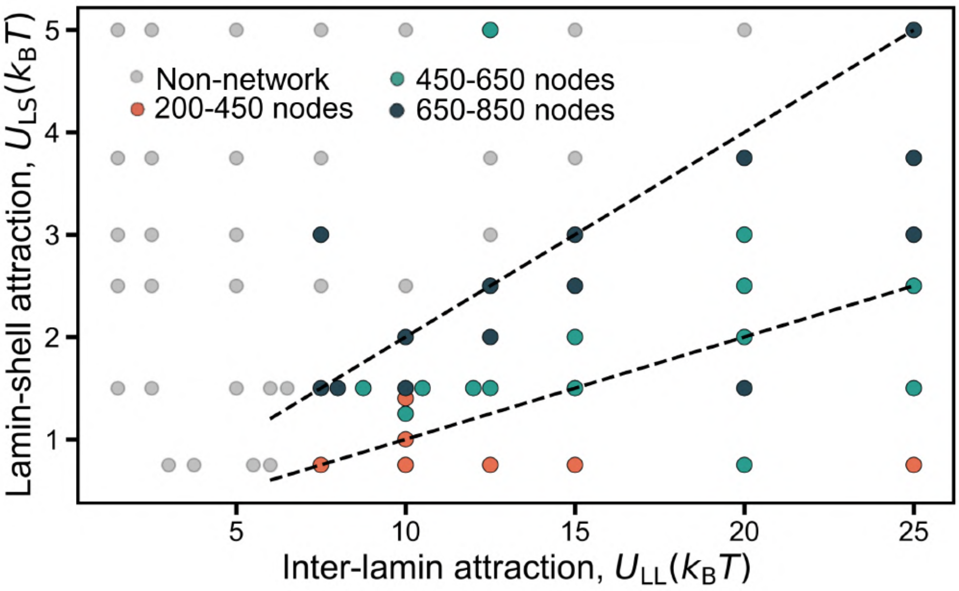


Figure S6: Average node count as a function of inter-lamin attraction, $U_{LL}$. Color codes represent i) gray: non-network cases, ii) blue and green: fibrous meshwork, and iii) orange: paracrystalline meshwork.

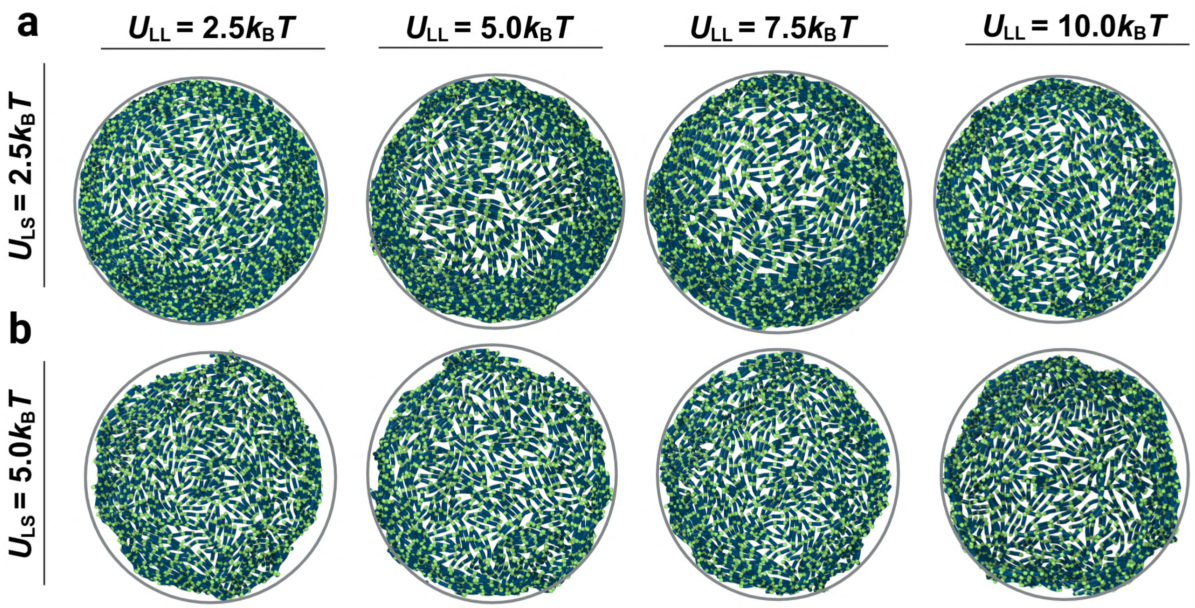


Figure S7: High lamin-shell ($U_{LS} \geq 2.5k_BT$) attraction leads to an isotropic phase irrespective of inter-lamin attraction, $U_{LL}$. (a) $U_{LS} = 2.5k_BT$ leads to an isotropic phase, while (b) $U_{LS} = 5.0k_BT$ leads to wrinkled shell shapes. Gray circles represent the spherical shell, provided there is no shape deformation.

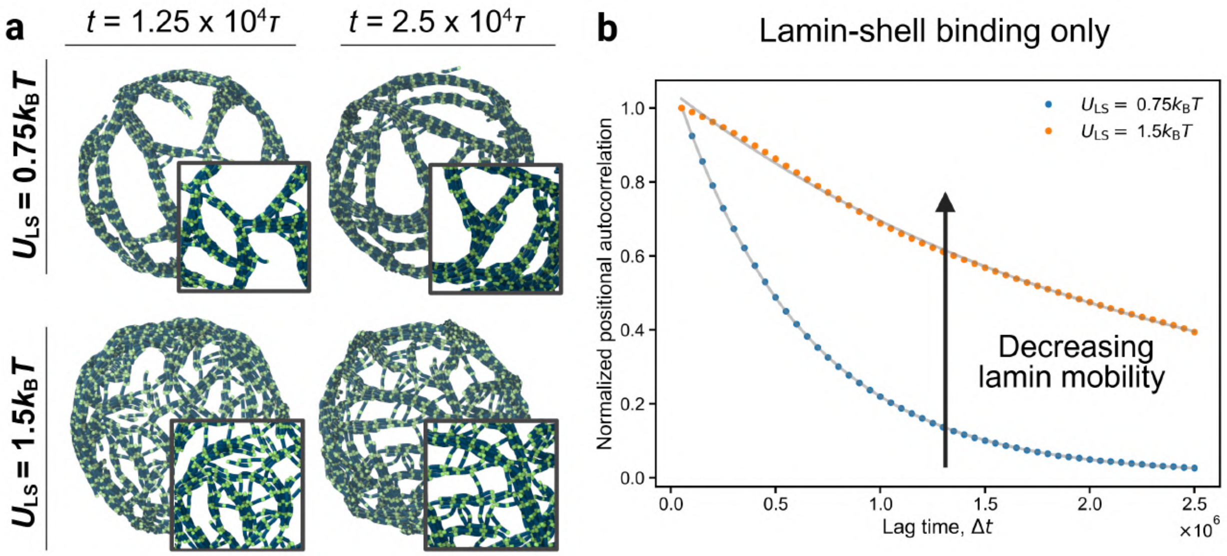


Figure S8: (a) Branched chains interconnecting paracrystalline arrays in a fibrous meshwork do not disappear upon doubling simulation time ($t = 1.25 \times 10^4\tau \rightarrow 2.5 \times 10^4\tau$). Inter-lamin attraction is fixed to $U_{\mathrm{LL}} = 10.0 k_{\mathrm{B}}T$. Thick, paracrystalline arrays remain intact when lamin-shell attraction is low, $U_{\mathrm{LS}} = 0.75 k_{\mathrm{B}}T$. Paracrystalline assemblies are connected via individual dimers when $U_{\mathrm{LS}} = 1.5 k_{\mathrm{B}}T$. (b) Increasing lamin-shell binding decreases lamin mobility. Normalized positional correlation function, $\langle r(t) \cdot r(t+\Delta t)\rangle / \langle r(t) \cdot r(t)\rangle$ was fitted to obtain decay times, $\tau$, and diffusion coefficients, $D$, shown in Table S2.

$$\frac{\langle r(t) \cdot r(t+\Delta t)\rangle}{\langle r(t) \cdot r(t)\rangle} = A \exp\left(-\frac{\Delta t}{\tau}\right) + C \tag{13}$$

where $A$ and $C$ are fitting constants, and the diffusion coefficient is calculated as

$$D = \frac{R^2}{2\tau} \tag{14}$$

where $R = 34\sigma$ is the radius of the elastic shell.

$$\frac{D_{0.75}}{D_{1.5}} = \frac{9.401 \times 10^{-4}}{2.676 \times 10^{-4}} = 3.51 \tag{15}$$

Stronger surface binding decreases chain mobility by approximately a factor of 3.5.

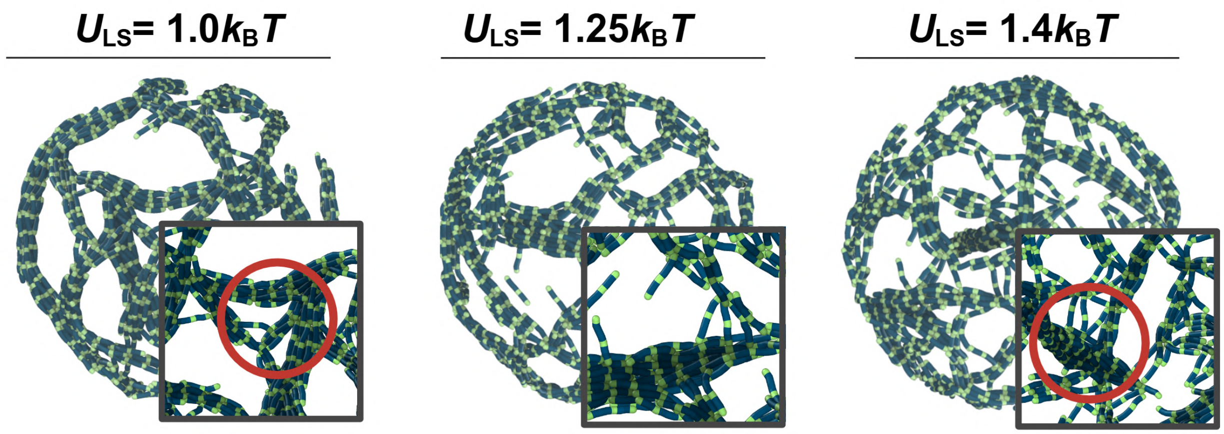


Figure S9: Probing lamin-shell attraction on the range $0.75 < U_{LS} < 1.5 k_B T$ reveals a transition between paracrystalline and fibrous meshworks driven by the mobility of lamin dimers on the surface (see main text). Red circles represent bulk aggregate formation.

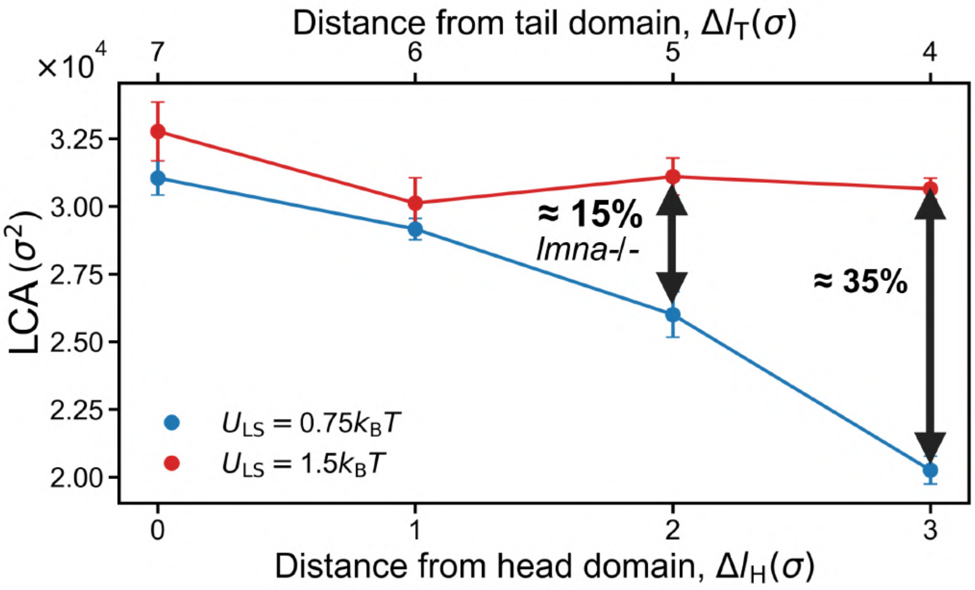


Figure S10: Lamin-covered area (LCA) as a function of distance from head domain, $\Delta l_H$ at the bottom, and distance from tail domain, $\Delta l_T$. We observe an inverse trend to face area, $A_{face}$ in the main text Fig. 3b. Percentages represent similar area alterations as upon removing lamin B [2].

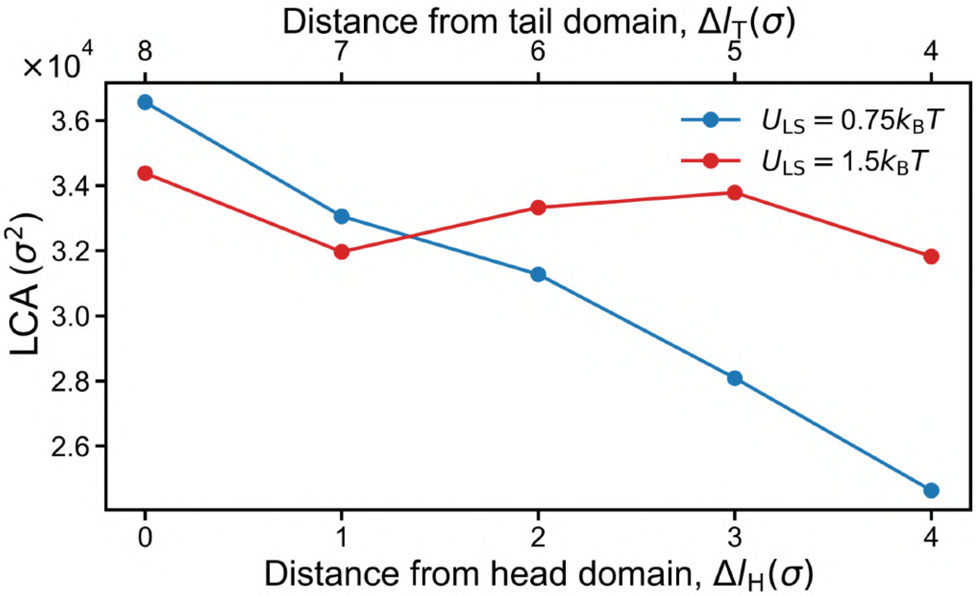


Figure S11: Lamin-covered area (LCA) as a function of distance from head domain, $\Delta l_H$ at the bottom, and distance from tail domain, $\Delta l_T$ as top axis when $n_{lamin} = 9\sigma$. We observe a similar trend as $n_{lamin} = 8\sigma$ lamin dimer in Fig. S19. The initial trend inversion is due to the presence of bulk aggregates marked in Fig. S13a.

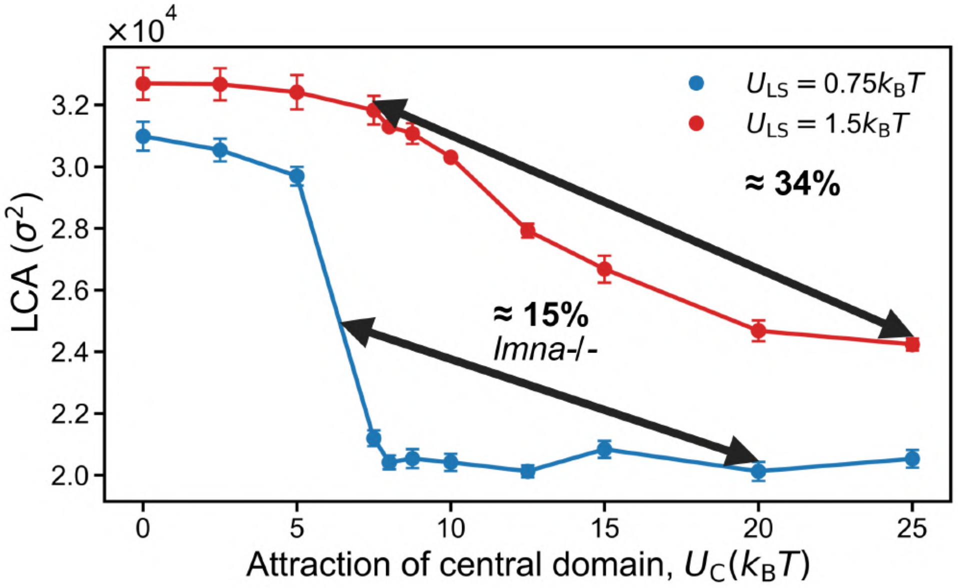


Figure S12: Lamin-covered area (LCA) as a function of central domain attraction, $U_C$.

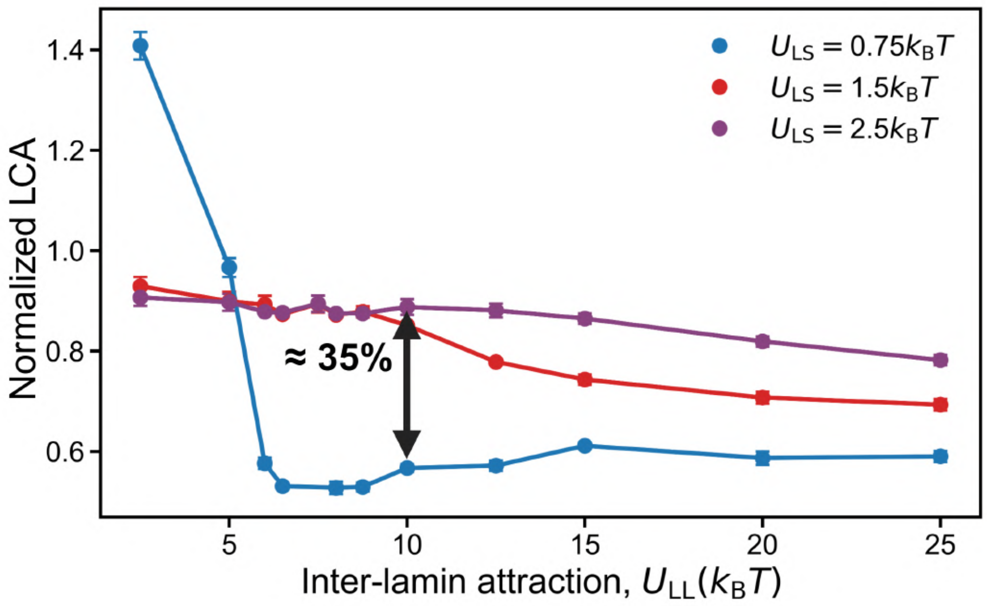


Figure S13: Normalized LCA decreases to < 1 with increasing inter-lamin attraction, $U_{\mathrm{LL}}$, analogous to trends observed in Fig. 3d-e.

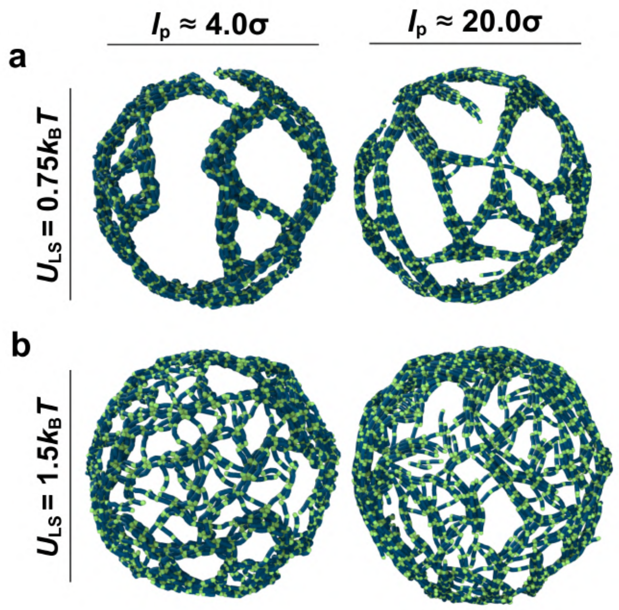


Figure S14: Lamin-shell attraction controls mesh opening size irrespective of stiffness of lamin dimers. Lamin dimers are made flexible by decreasing their persistence length to $l_{\mathrm{p}} = 4.0\sigma$ such that $l_{\mathrm{p}}/l$ decreases from 2.5 (default and right column) → 0.5 (left column).

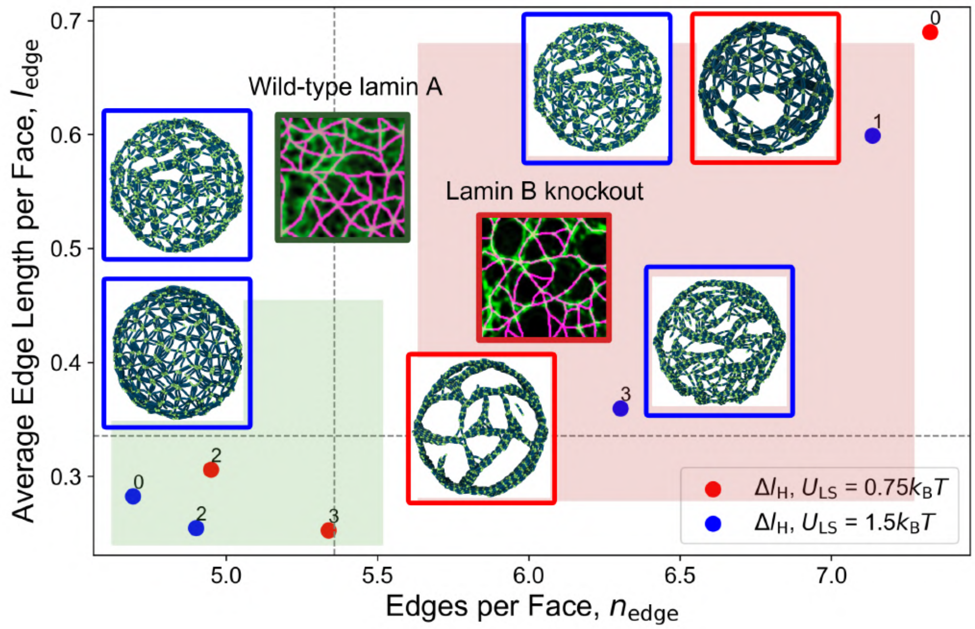


Figure S15: Effect of the distance of additional attractive domain, $C$, from the head domain, $\Delta l_{\mathrm{H}}$, on lamina meshwork properties. A central attractive site affects both $l_{\mathrm{edge}}$ and $n_{\mathrm{edge}}$, similar to *lmnb1*-/- nuclei [2]. No central attraction site leads to non-homogeneous face sizes [2, 10, 11].

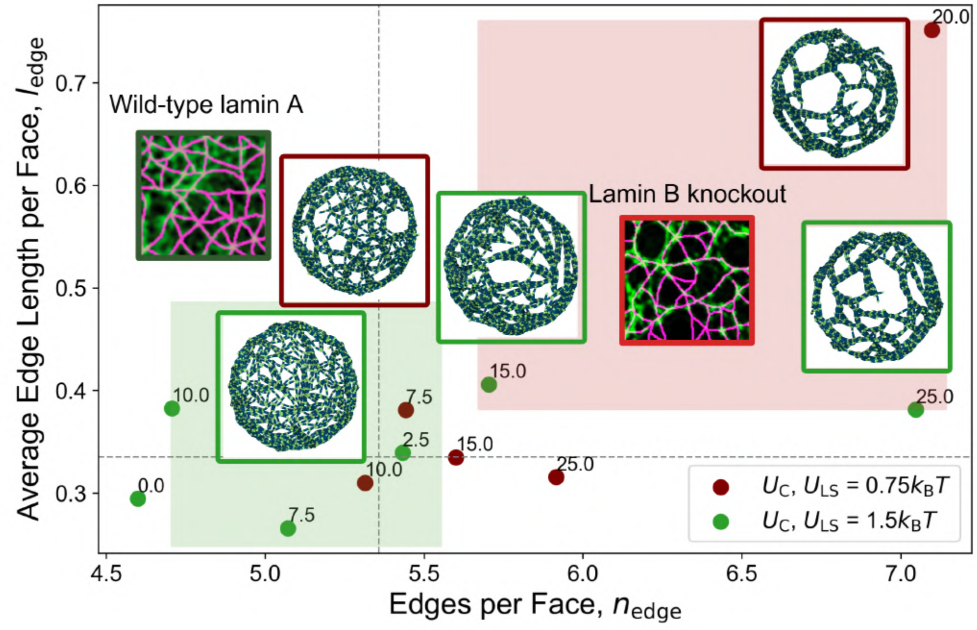


Figure S16: Effect of the strength of the central attraction, $U_{\mathrm{C}}$, on on lamina meshwork properties. When $U_{\mathrm{C}} \gg U_{\mathrm{LL}} = 10.0 k_{\mathrm{B}}T$, we observe continuous paracrystalline arrays, which might characterize meshworks with large opening sizes [2, 12].

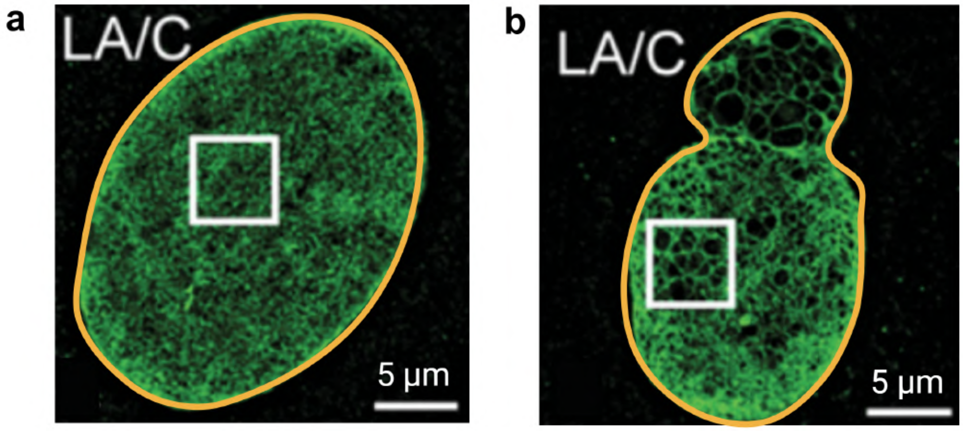


Figure S17: (a) Wild-type and (b) lamin B knockout nuclei used to quantify nuclear surface area, $A_{\text{surf}}$ shown in Table S4. Yellow outlines represent the nuclear area taken into account. Scale bar: 5 $\mu m$. Images are obtained from [2].

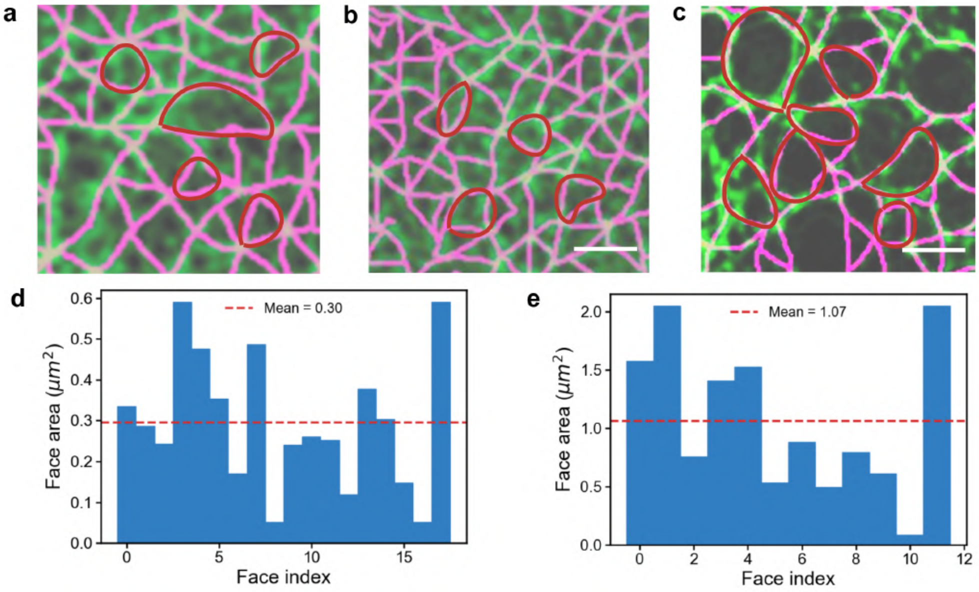


Figure S18: 3D-SIM images of (a-b) wild-type and (c) lamin B knockout lamina used to quantify experimental face area, $A_{\text{face}}$ shown in Table S4. Red outlines represent how the face area is measured. Scale bar: 1 $\mu m$. Images are obtained from [2]. Experimental face area distributions for (a) wild-type and (b) lamin-B knockout nuclei.

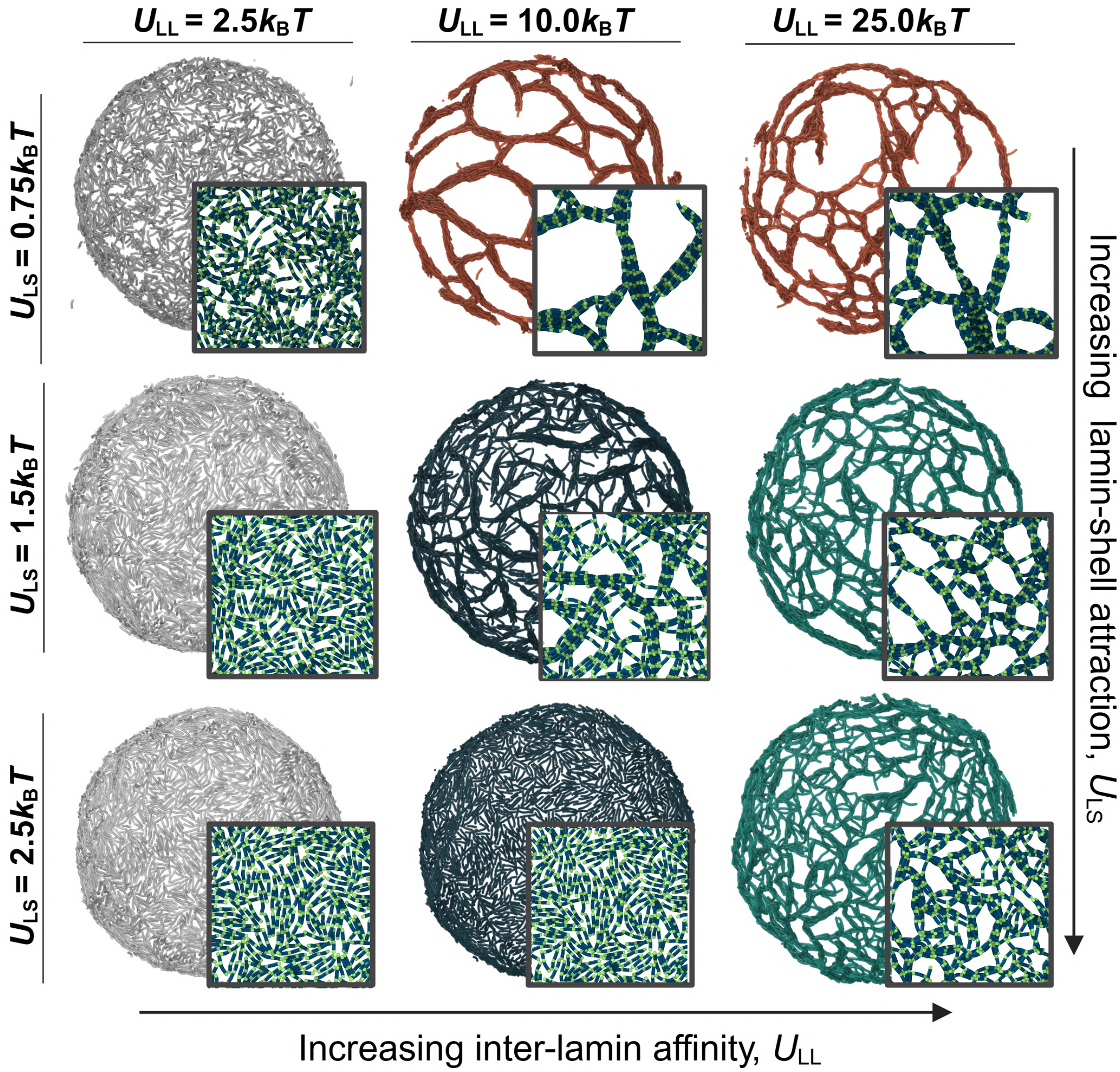


Figure S19: Increasing the system size yields the same trends as shown in Fig. 3b, provided that the surface coverage remains constant. The system size is doubled by increasing the elastic shell radius from $R = 34\sigma$ to $R = 68\sigma$, which reduces the lamin length-to-shell radius ratio from $l/R = 0.25$ to 0.1. The surface concentration is kept constant and calculated as

$$c_{\text{surface}} = \frac{N_{\text{lamin}} \cdot n_{\text{lamin}}}{A} = \frac{1250 \cdot 8}{4\pi(34\sigma)^2} = \frac{5000 \cdot 8}{4\pi(68\sigma)^2} \approx 0.69\,\sigma^{-2}$$

where $A$ is the surface area occupied by lamin dimers.

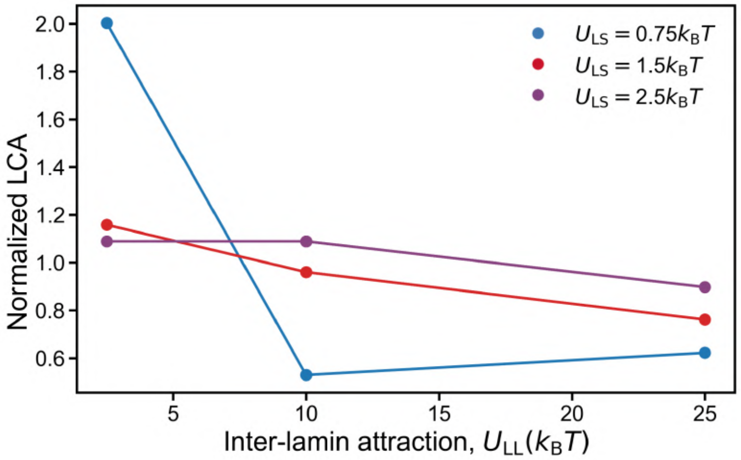


Figure S20: Normalized LCA exhibits similar trends with lamin interactions, irrespective of elastic shell radius and system size (compare to Fig. S22 and Table S4).

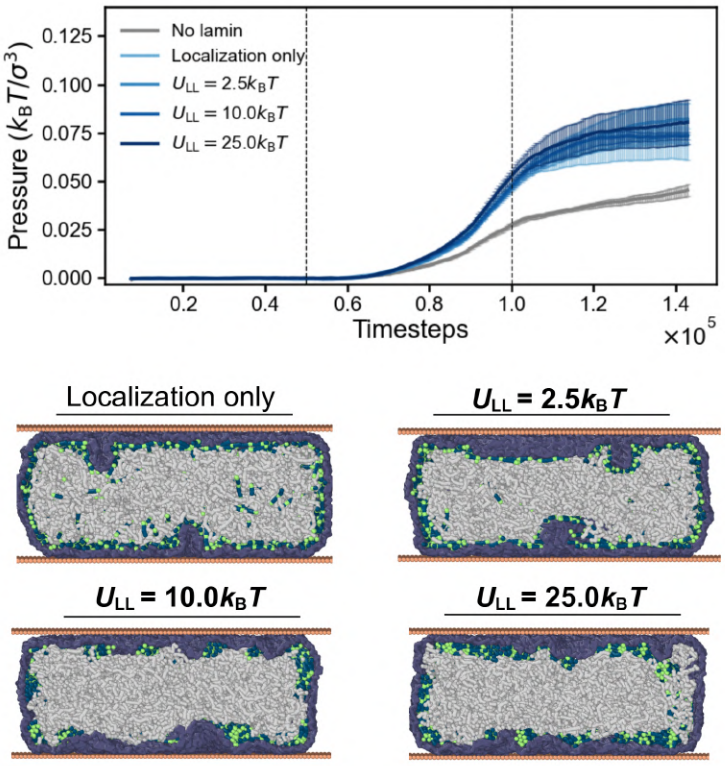


Figure S21: Effect of inter-lamin attraction on stress-time curves when lamin-shell binding is weak, $U_{LS} = 0.75 k_B T$. Snapshots show shell deformation after compression for corresponding cases.

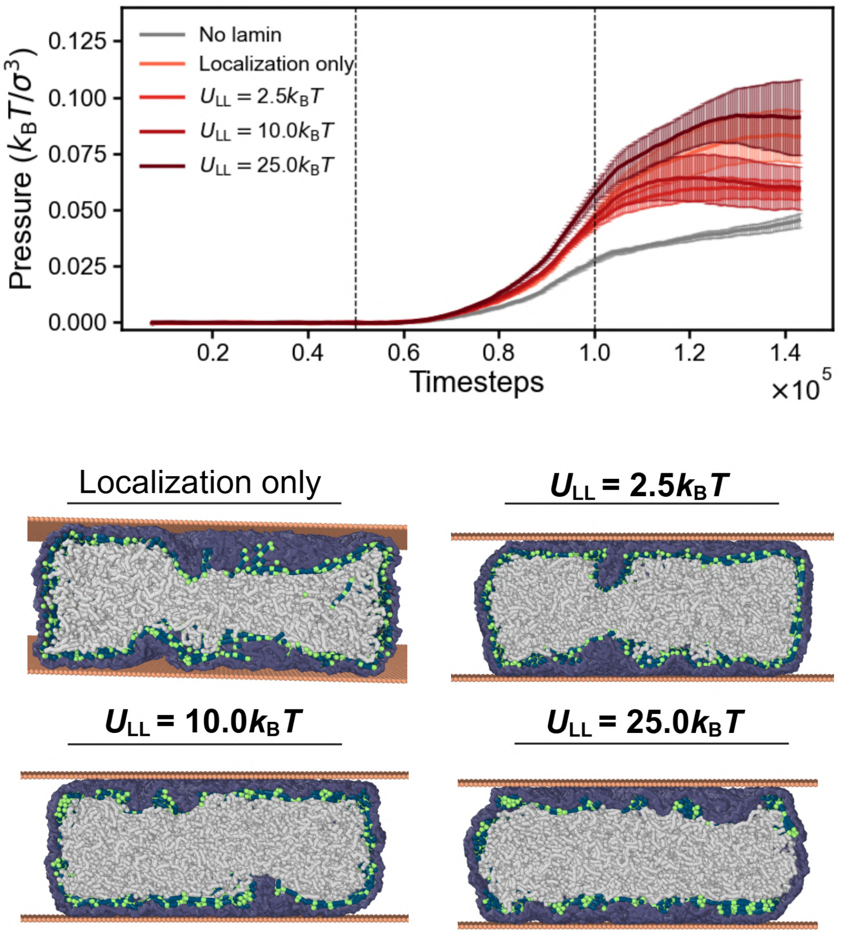


Figure S22: Effect of inter-lamin attraction on stress-time curves when lamin-shell binding is strong, $U_{LS} = 1.5k_BT$. Snapshots show shell deformation after compression for corresponding cases.

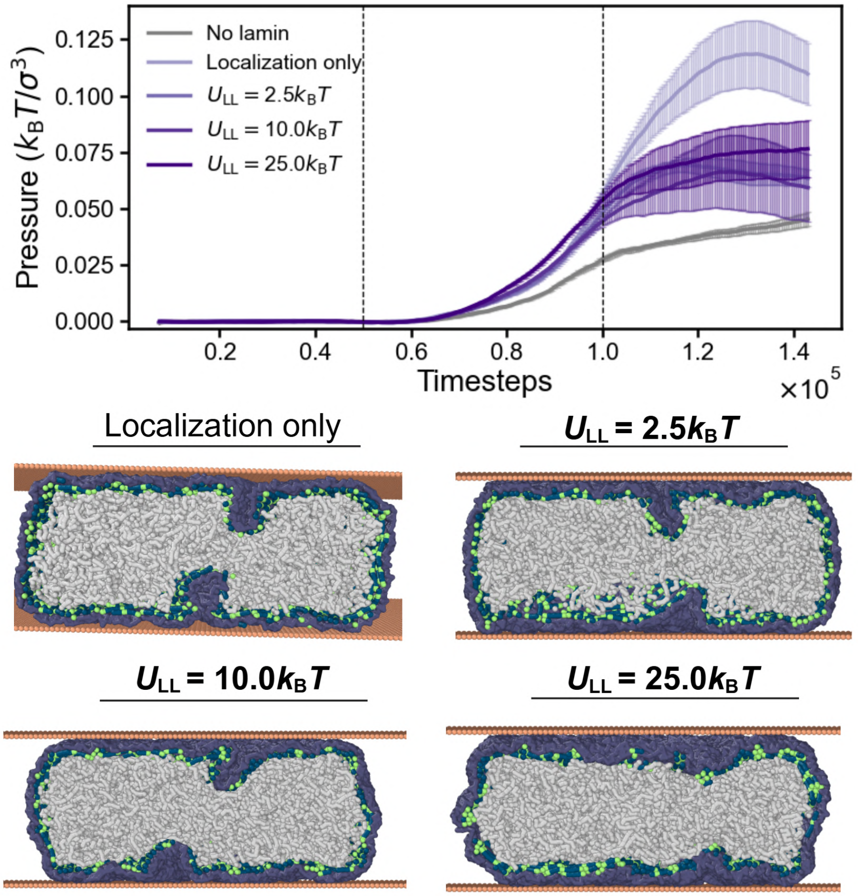


Figure S23: Effect of inter-lamin attraction on stress-time curves when lamin-shell binding is strong enough to wrinkle the shell, $U_{LS} = 2.5k_B T$. Snapshots show shell deformation after compression for corresponding cases.

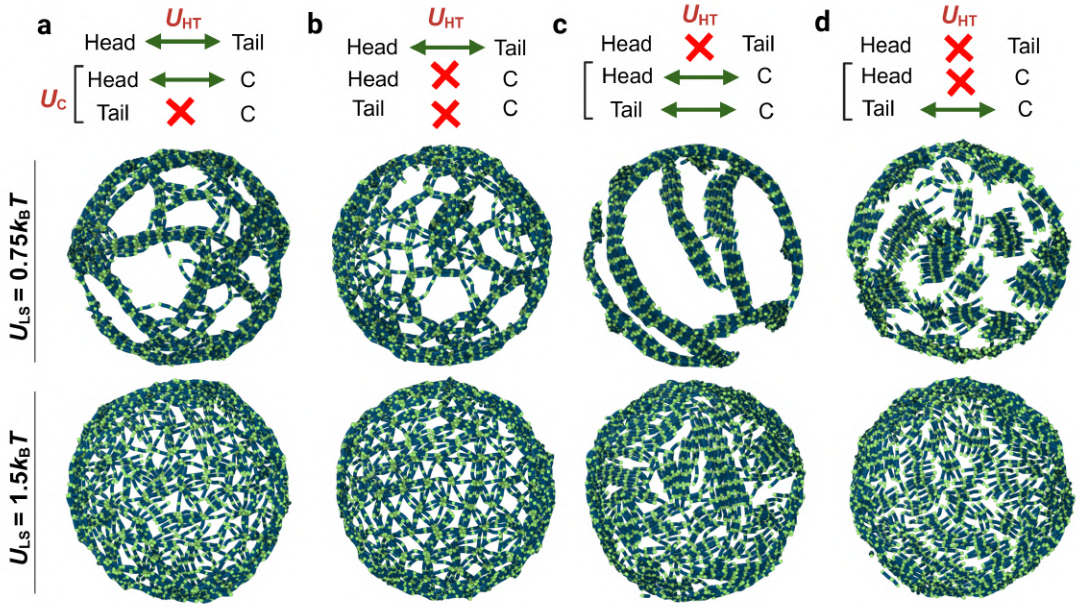


Figure S24: Schematic and simulation snapshots of perturbations to inter-lamin interactions with inter-lamin attraction fixed to, $U_{LL} = 10.0k_B T$ ($U_{LL} = 25.0k_B T$ in main text Fig. 4d-g).

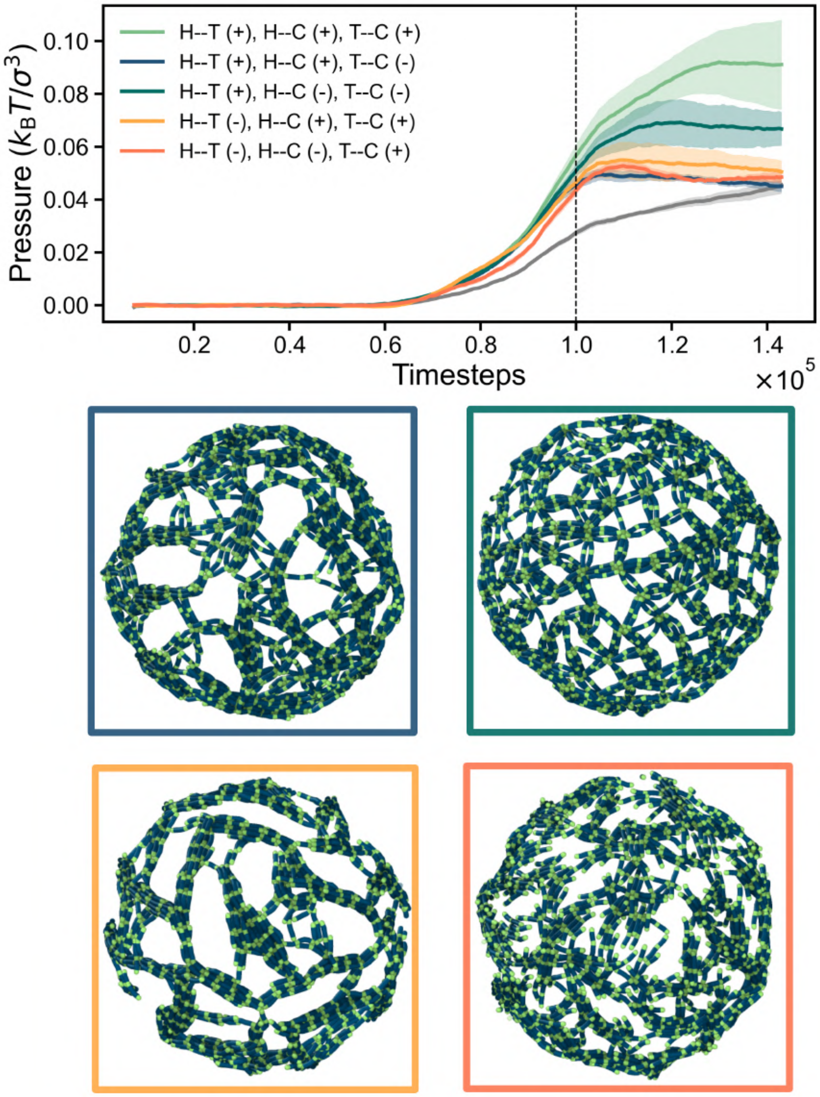


Figure S25: Pressure-time curve for mutations under strong lamin-shell binding, $U_{\mathrm{LS}} = 1.5 k_{\mathrm{B}} T$ (see main text, Fig. 4g, right panel).

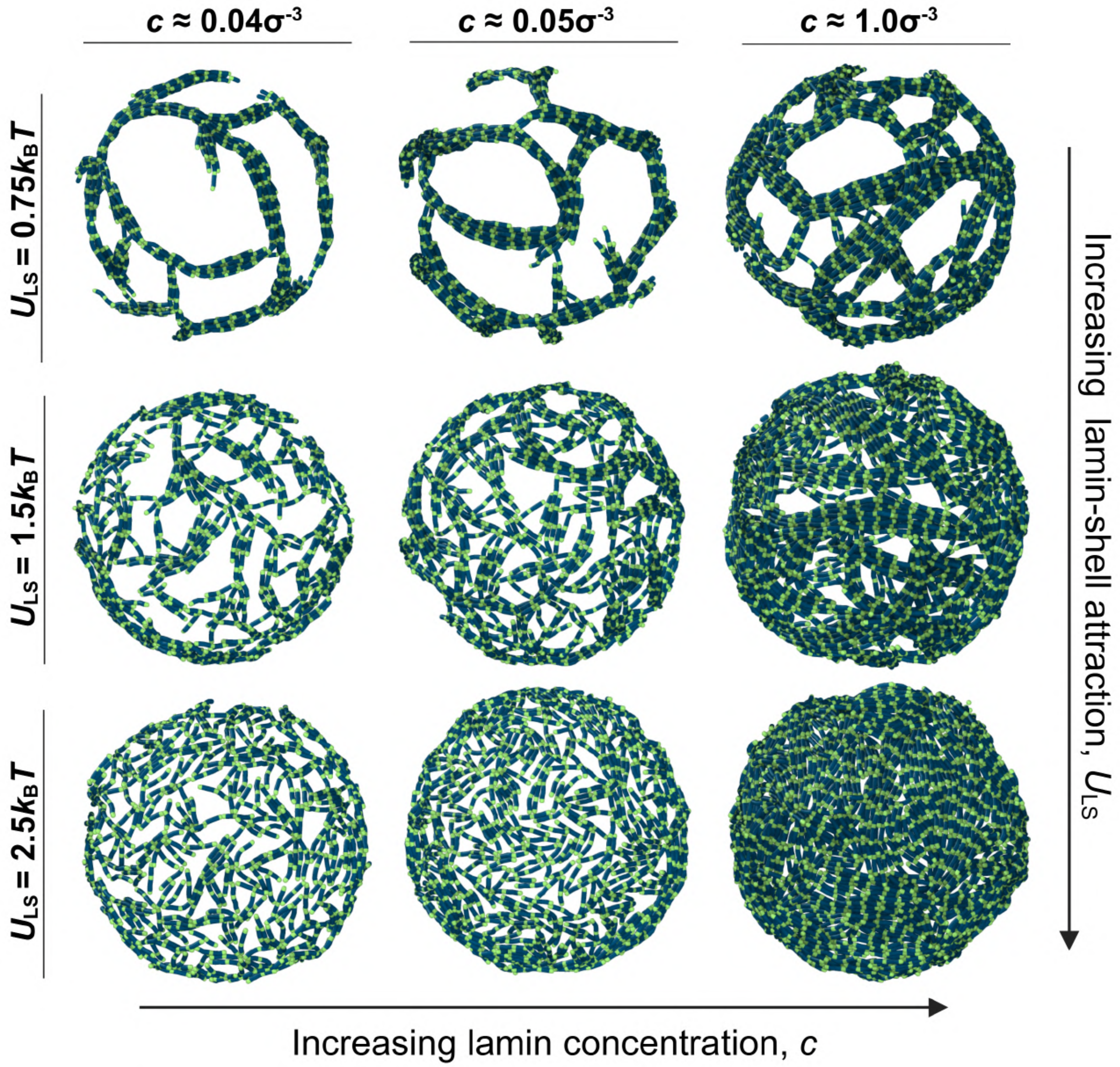


Figure S26: Effect of lamin concentration on paracrystalline array formation.

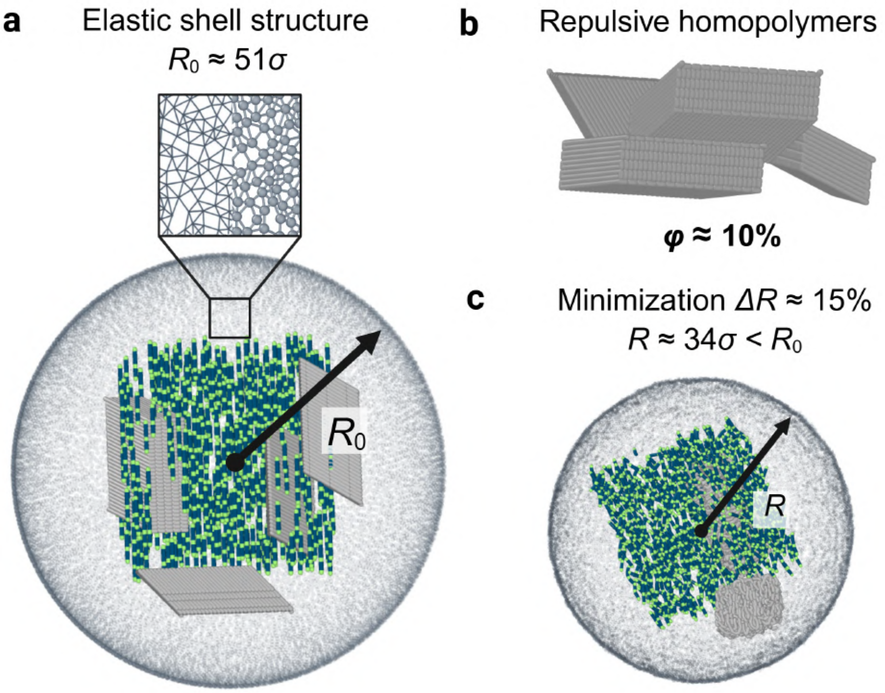


Figure S27: Simulation setup for the MD lamina model. (a) Bonded elastic shell with lamin dimers and homopolymers at the interior. (b) $n_{\text{polymer}} = 4$ repulsive homopolymers model the osmotic pressure of the genome at the shell interior. (c) The initial structure in (a) is minimized for $30\tau$ using an integration time step of $\Delta t = 0.001\tau$. This step ensures that the shell is tightened before the interior contents mix [9].

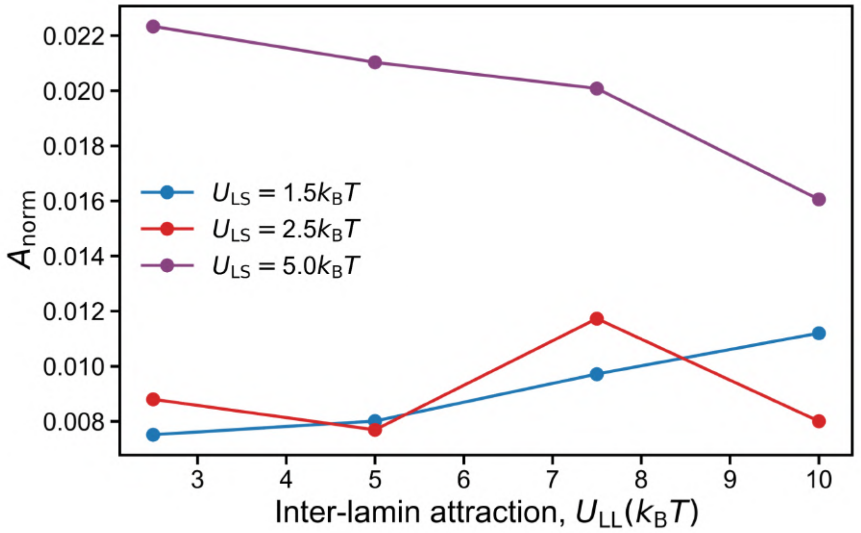


Figure S28: Normalized asphericity, $A_{\text{norm}}$, increases by an order of magnitude when lamin-shell attraction is increased to $U_{\text{LS}} = 5.0k_{\text{B}}T$, indicating moderate shell shape distortion.

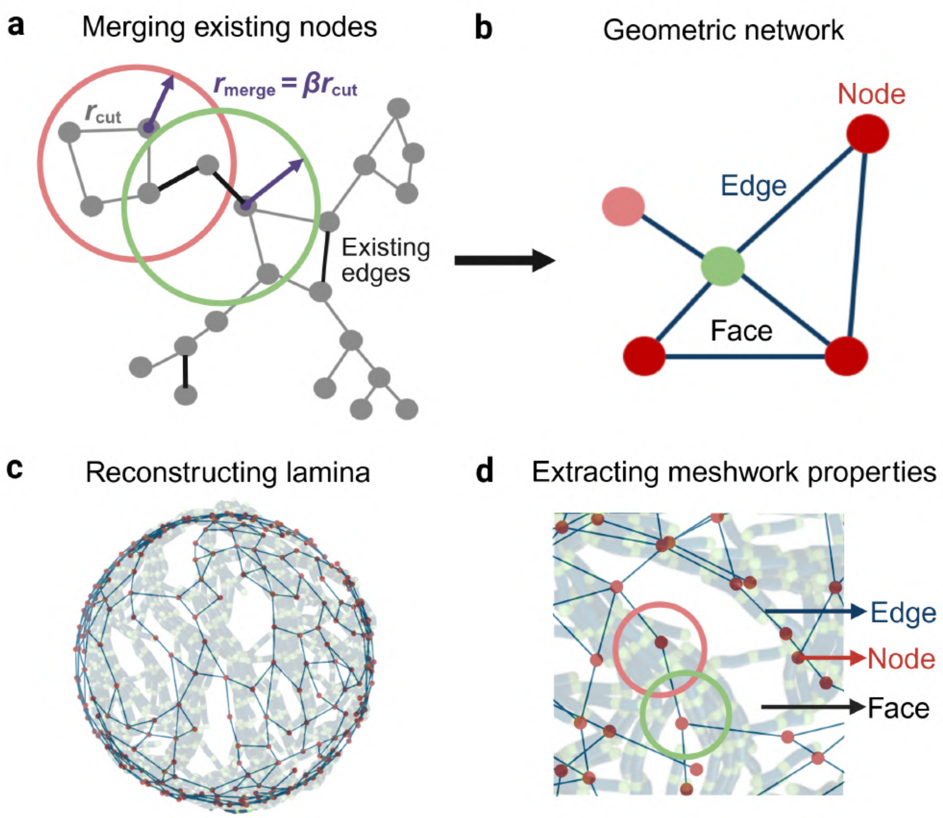


Figure S29: Network reconstruction methodology to quantify lamina meshwork properties. Schematic representation of (a) lamina meshwork and its (b) reduced lamina network after position-based node merging. (c) 3D reconstructed geometric network, a single lamina simulation. (d) Metrics we can obtain from this methodology.

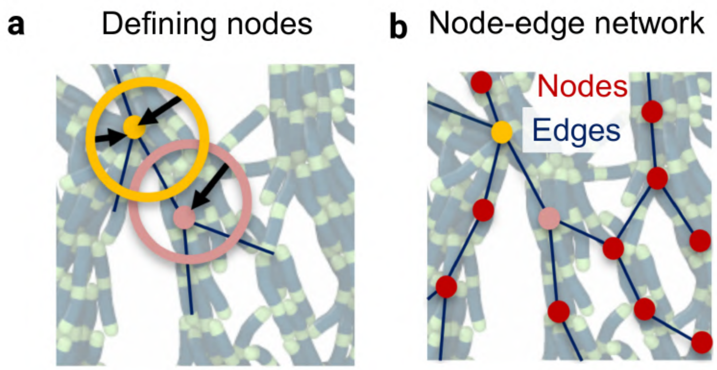


Figure S30: Simplified representation of Fig. S3 representing the (a) node clustering algorithm, and (b) corresponding node-edge representation.

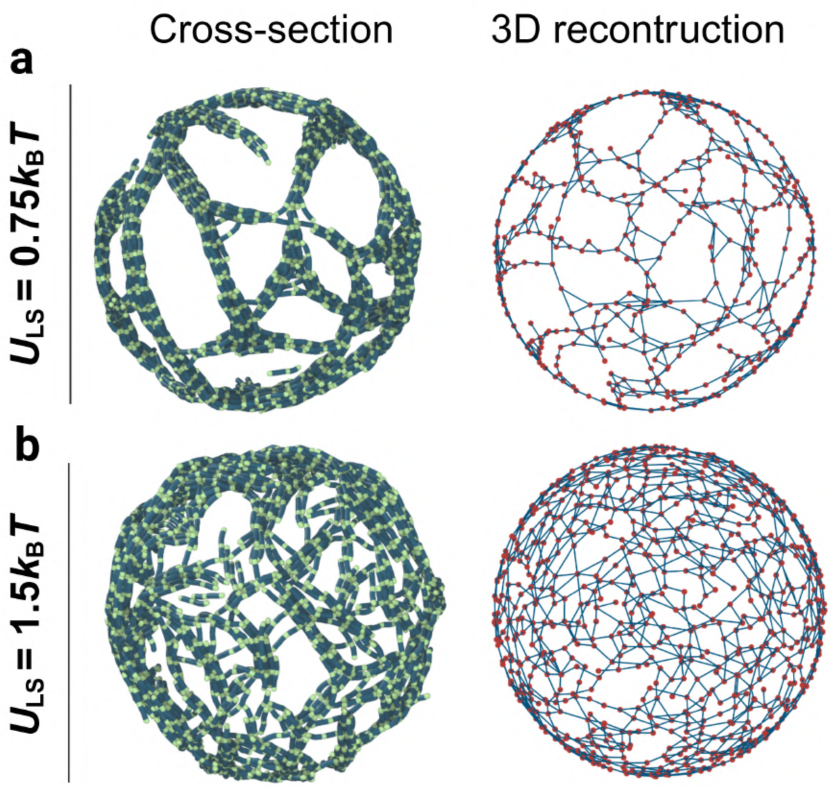


Figure S31: Reconstructed network for (a) paracrystalline and (b) fibrous meshwork.

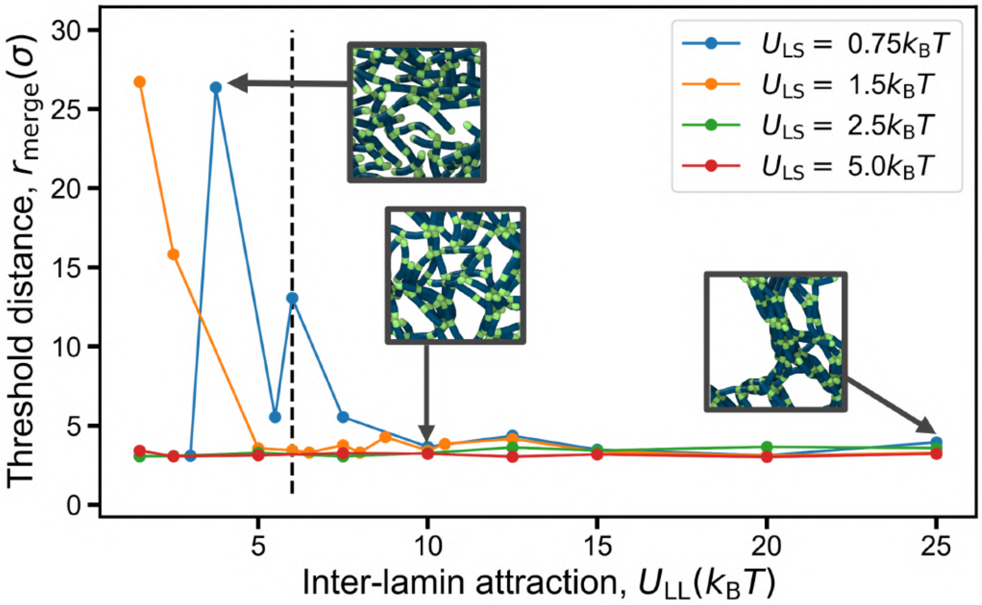


Figure S32: Network reconstruction for various isotropic arrangements ($U_{LL} \leq 6.0 k_B T$) requires very high thresholds, $r_{merge} > 5.5\sigma$.

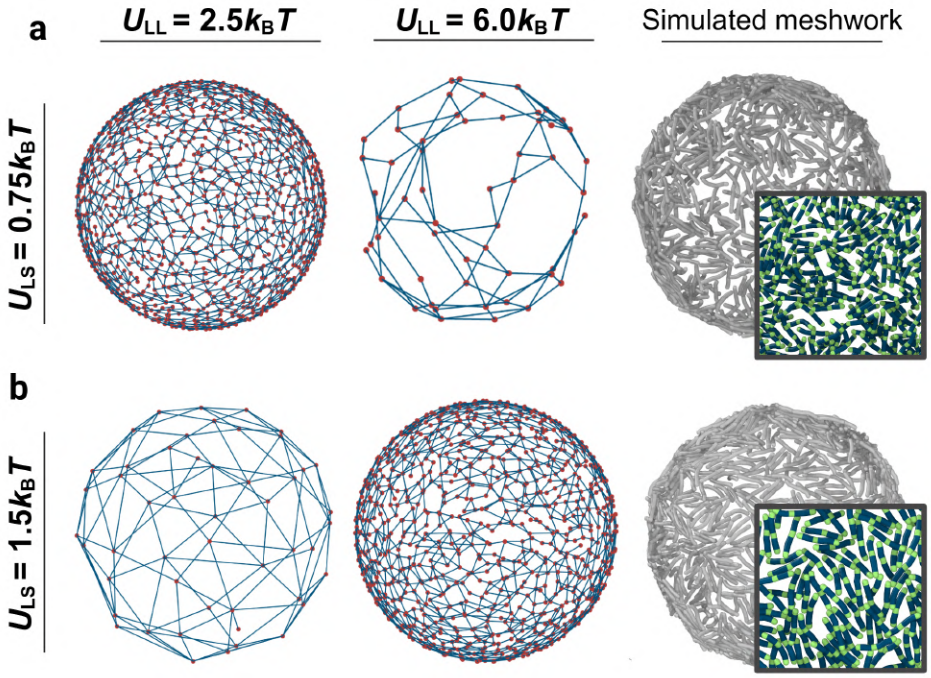


Figure S33: Reconstructed meshworks for non-network cases, when $r_{\text{merge}} > 5.5\sigma$. 3D networks are either over-connected (average node count ≥ 850) or highly disconnected (average node count ≤ 150) compared to the meshwork arrangement.

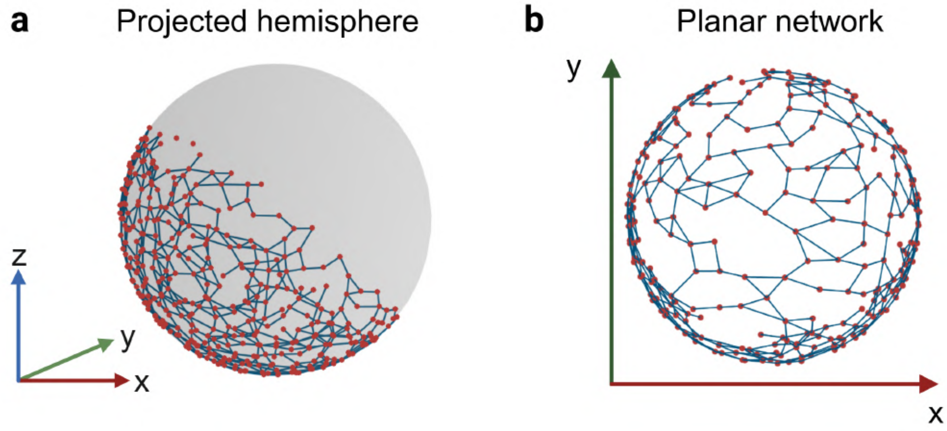


Figure S34: Number of faces of the reconstructed network is quantified by reducing the 3D network (a) to a slice in the $xy$ plane along the center (b).

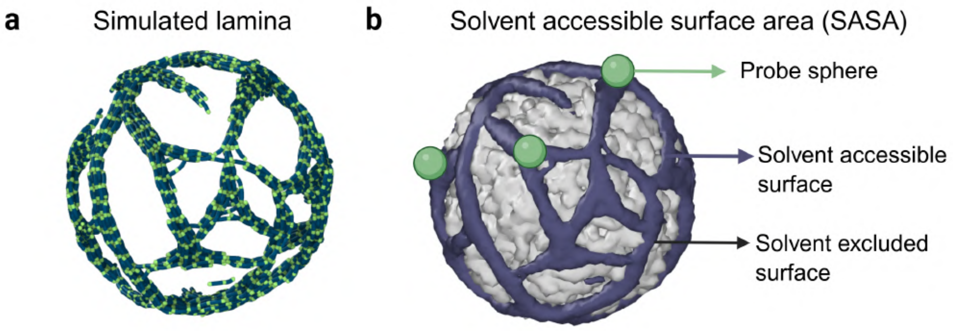


Figure S35: Schematic representation of how the lamin-covered area (LCA) is calculated from solvent-accessible surface area (SASA). (a) Cross-section of simulated lamina. (b) Spheres of radii of 0.5$\sigma$ probe the meshwork surface (purple), yielding an effective surface area. Regions not covered by lamin fibers make up the solvent-excluded surface (gray).

### 6 Supplementary Movies

- Movie S1: Simulation movie of weak lamin binding, $U_{\mathrm{LS}} = 0.75 k_{\mathrm{B}} T$, during the relaxation phase after compression.
- Movie S2: Simulation movie of weak lamin binding, $U_{\mathrm{LS}} = 2.5 k_{\mathrm{B}} T$, during the relaxation phase after compression.
- Movie S3: Movie of uniaxial compression simulation.
